# A systematic examination of learning in the invasive ant *Linepithema humile* reveals very rapid development of short and long-term memory

**DOI:** 10.1101/2022.04.12.487867

**Authors:** Thomas Wagner, Henrique Galante, Roxana Josens, Tomer J. Czaczkes

## Abstract

The Argentine ant (*Linepithema humile*) is one of the most damaging and widespread invasive ant species worldwide. However, control attempts often fail due to bait rejection or insufficient bait uptake. Increasing preference for, and consumption of, bait is thus an important requirement for successful control. Learning and within-nest information transfer might be a potential tool for achieving this goal. We conducted a systematic investigation of olfactory learning and route learning in Argentine ants. The ants showed very strong and rapid route learning, choosing the correct arm in a Y-maze 65% of time after just one visit, and 84% correct after two. Odour learning was even more rapid, reaching up to 85% correct choices after just one exposure to flavoured food. Learning is long-lasting, with 73% correct choices after 48h. Food flavour information is transferred efficiently between nestmates in the nest, driving preference: naïve ants housed with ants fed on flavoured food show a strong preference (77%) for that odour after 24h. Overall, *Linepithema humile* are outstanding learners. This, coupled with efficient intranidal information transfer and strong pheromonal recruitment, may help explain their ability to discover and then dominate resources. However, these strengths could potentially be used against them, by exploiting learning and information transfer to increase toxic bait uptake. Steering ant preference by leveraging learning might be an underappreciated tool in invasive alien species control.

## Introduction

With increasing trade, and the concomitant breakdown of biogeographic barriers, invasive species are spreading globally (Mack and Lonsdale 2001; Mooney and Cleland 2001; Perrings et al. 2005; Bates et al. 2020). Invasive species are often economically damaging and ecologically devastating (Marbuah et al. 2014; Escobar et al. 2018; Angulo et al. 2022), with invertebrates being amongst the most damaging invasive groups (Ricciardi 2015). Ants are especially effective at invading habitats outside their native range (Tsutsui and Suarez 2003), where they displace native species through competition and predation (Ness and Bronstein 2004; Abbott 2006; Matsui et al. 2009).

The Argentine ant (*Linepithema humile* (Mayr, 1868)) is one of the most damaging ant species worldwide, and the most widespread invasive ant in Europe (Human and Gordon 1999; Trigos-Peral et al. 2021; Angulo et al. 2022). The presence of *L. humile* in invaded areas causes a massive decrease in invertebrate diversity and even affects vertebrates (Wetterer et al. 2001; Suarez et al. 2005; Alvarez-Blanco et al. 2021). In addition, *L. humile* can act as an important agriculture pest by enhancing Hemipteran populations, which then increase the likelihood of fungal and viral infections (Wetterer et al. 2009).

Combatting invasive ants has become a top priority for conservation programs (Jenkins and Forte 1973; Rust et al. 2003; Sunamura et al. 2011; Hoffmann et al. 2016; Sakamoto et al. 2019). Unfortunately, in addition to being ecologically and economically damaging, invasive ants are also difficult to control. Two-thirds of *L. humile* eradication attempts have failed (Hoffmann et al. 2016). Insect control methods usually rely on the use of insecticide spraying. However, the success rate of this technique against ants is known to be highly depanding on various factors and circumstances such as weather condition, landscape surface, colony density, and control effort (Jenkins and Forte 1973; Bhandari et al. 1997; Klotz et al. 2007). A major reason for the often poor success of insecticide spraying on ants is that the queens and brood are physically sheltered (Williams et al. 2001; Rust et al. 2003). A currently promising approach for eradication of ants is to use baits with a slow-acting poison, which workers return to the brood and the queen (Hoffmann et al. 2016). Even so, the success rate of such eradication attempts has been low (Souza et al. 2008; Hoffmann 2011). A big issue is the availability of high-quality natural food, which is often preferred by ants and acts as a competitor for the poisoned bait, leading to low bait consumption rates (Rust et al. 2003; Silverman and Brightwell 2008). Driving bait preference and increasing consumption is thus a critical step towards successful ant control.

One approach to increase bait attractiveness, beyond a change in bait formulation, is to steer ant behaviour. Studies in grass-cutting ants (Hughes et al. 2002) and leaf-cutting ants (Knapp 1987, 1995) suggested that alarm pheromones may be used to increase bait consumption and foraging activity. A study in Argentine ants showed that synthetic (Z)-9-hexadecenal, an aggregation factor, increased bait (liquid sucrose) consumption (Greenberg and Klotz 2000) and the mortality rate in an insecticide spray usage under laboratory conditions (Choe et al. 2014). Pheromones can not only be used to lure ants to a bait, but also to disrupt their trail-following behaviour (Tatsuki et al. 2005; Tanaka et al. 2009). An open field experiment demonstrated that the combination of a highly-concentrated synthetic pheromone and insecticidal baits may provide effective control of Argentine ant populations (Sunamura et al. 2011).

However, preference and consumption could also potentially be manipulated by exposing individuals to tailored information. Possibly the easiest way of changing the perceived value of a cue is via associative learning. For example, bees which are fed on sunflower nectar scented food in the nest preferentially visit sunflower fields (Farina et al. 2020a). This demonstrates that training based on associative learning can steer individual and collective insect preference. It may be possible to likewise steer the preference of invasive and pest species towards baits and away from competing natural food sources. However, a key prerequesit for this is strong associative learning abilities in the target species.

Ants have been shown to be excellent learners. They can form complex route memories which allow them to associate a direction, panorama, or a route with a food reward, nest (Aron et al. 1988; Harrison et al. 1989; Graham and Collett 2006; Grüter et al. 2011; Knaden and Graham 2016) or even with a negative outcome (i.e. getting trapped) (Wystrach et al. 2020). Many ant species have been shown to associate an odour with a food reward (Roces 1990; Helmy and Jander 2003; Dupuy et al. 2006; Czaczkes et al. 2014; Oberhauser et al. 2019; Czaczkes and Kumar 2020). Food-odour associations can be rapidly formed, sometimes after only one exposure, and may last for days (Josens et al. 2009; Arenas and Roces 2018; Piqueret et al. 2019).

However, very little is known about the associative learning ability of invasive ants. In one study it was shown that Argentine ants (*L. humile*) can use visual and spatial cues to find a food source, but the experimental setting was a binary choice test between visual cues and a pheromone trail (Aron et al. 1988, 1993). A recent study showed that they are also able to associate an odour spot in a circular arena with a sucrose solution (Rossi et al. 2020). Preexposure to trail pheromone increased food acceptance for low concentration sucrose solutions, but had no effect on associative learning. Rossi et al. (2020) is a very important paper for us, as it is the only currently available investigation of associative learning in free-running Argentine ants. However, it is unclear how many exposures to the reward would be required in order for the individual to form a short-term association. Furthermore, no long-term memory tests were conducted, nor has information transfer of food-associated odours between nestmates been investigated, as previously shown in bees, wasps, and other ants (Farina et al. 2005; Provecho and Josens 2009; Schueller et al. 2010).

Here, we conduct a comprehensive investigation of learning in the ecologically important ant *L. humile*. We study how rapidly *L. humile* form short- and long-term memories, which types of cues best support this, and whether food-related cues are transferred between nestmates intranidally. Such information is critical if we hope to develop cognition-based control strategies.

## Materials and Methods

### Colony maintenance

*Linepithema humile* ants were collected in 2021 from Girona, Spain and Proença-a-Nova, Portugal, and were all part of the same European supercolony. Colony fragments (henceforth colonies), consisting of one or more queens and 300-1000 workers, were kept in plastic foraging boxes (32.5 × 22.2 × 11.4cm) with plaster of Paris on the bottom. The walls were coated in fluon to prevent escape. Each box contained several 15mL plastic tubes, covered in red foil, and partly filled with water and plugged with cotton, for use as nests. The ants were maintained on a 12:12 light:dark cycle at room temperature (21-25 °C) and provided with water *ad libitum*. Colonies were fed for three days with *ad libitum* 0.5M sucrose solution and freeze-killed *Drosophila melanogaster*, and deprived of sucrose and *Drosophila* for four days prior to testing. For all experiments using odours and/or flavours, donor and recipient colony pairs were used. Such colonies were collected in the same location at the same time. However the donor colonies had never experienced any of the odours and flavours used (see below), whilst the recipient colonies received individuals that had experienced these. All experiments were conducted between June and September 2021 except the 6^th^ and the 7^th^ experiment (December 2021) and the long-term experiments 8 and 9 (October 2021)

### Solutions and odours

1M sucrose solutions (Südzucker AG, Mannheim, Germany), were used as a reward during training for all experiments. Where a negative reinforcement was also presented, 0.6mM quinine (Merck KGaA, Darmstadt, Germany) solution was used. Scented paper overlays, used during experiments to provide environmental odours, were stored for at least 1 week prior to the experiments in airtight plastic boxes (19.4 × 13.8 × 6.6cm) containing a glass petri-dish with 500μL of either strawberry or apple food flavouring (Seeger, Springe, Germany). For experiments where flavoured food was used, 1μL of the respective flavouring was added per 1mL of 1M sucrose solution. Pilot studies showed *L. humile* workers having a slight preference for strawberry over apple odour (58% of the ants prefered strawberry, N = 158, see supplement 3).

### Route association: Short-term spatial-memory

#### Experiment 1 – Short-term spatial-memory

Here we investigate side learning by offering sucrose on one arm of a Y-maze and bitter quinine on the opposing arm. 6-8 ants were tested per day, testing in total 46 ants from 7 colonies.

Y-maze setup and training methods follow Czaczkes (2018): A colony was connected via a drawbridge to a Y-maze (arms 10cm long, 1cm wide, tapering to 2mm at the bifurcation, see fig. 1a) covered in unscented disposable paper overlays. A c. 20μL drop of sucrose solution was placed at the end of one arm of the maze, and a drop of quinine solution on the other. The first 1-3 ants to initially choose the arm leading to the quinine-laden arm were marked with acrylic paint after they subsequently found the sucrose reward. This protocol ensures that the focal ants do not have an innate preference for the rewarded side. From this point on, only the marked ants were selectively allowed to move onto the setup, with only one ant allowed onto the maze at a time. Upon satiation, ants ran back over the bridge to the nest and unloaded their collected food to their nestmates. While unloading, the paper overlays were replaced with fresh overlays, to remove any pheromone trails or cuticular hydrocarbons left by the ant. After unloading, the ants were allowed back onto the Y-maze. We recorded the ant’s initial decision (defined as the antennae crossing a line 2cm from the bifurcation) and final decision (crossing a line 8cm from the bifurcation, 2cm from the arm end). We then allowed the ants to carry out 3 more such visits (5 visits to the sucrose, 4 trained decisions in total). Reward position was systematically varied between ants. The Y-maze was cleaned with ethanol after each group of 1-3 ants was finished.

**Figure 1:**
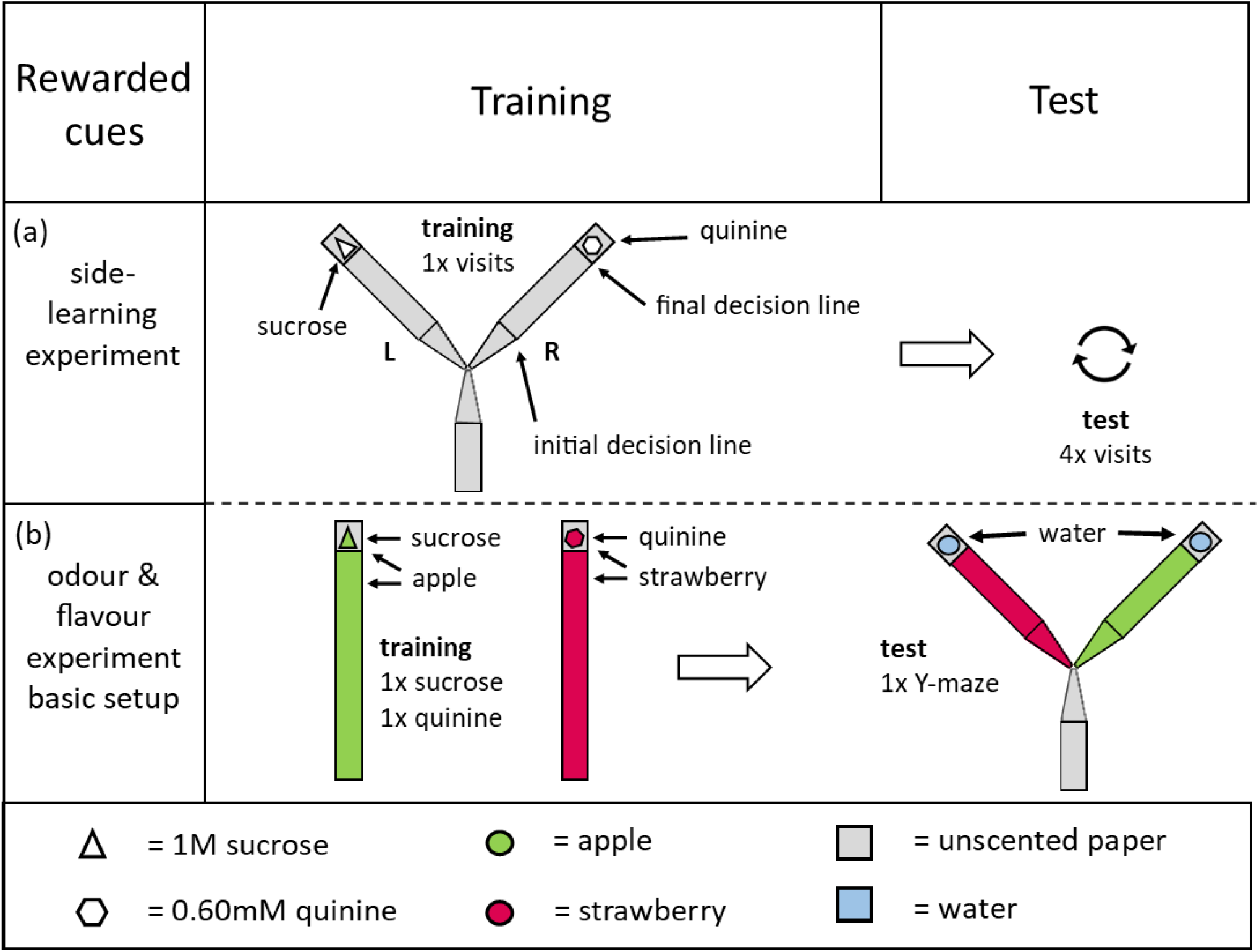
Basic setup used for all experiments: a) In the side-learning experiment, ants were trained to associate a side of the Y-maze with a 1M sucrose reward, while the other side contained 0.60mM quinine or water (exp 1). In all other experiments (b), ants were trained to associate an odour and/or a flavour (apple or strawberry) with a 1M sucrose reward on a linear runway, and were then tested on a Y-maze. In experiments where quinine was used as negative reinforcer (2,4,6,7) the flavour and odour of the quinine was the converse of that of the reward. In the experiments (3,5,8,9) the quinine punishment training visit was omitted. In the test trials the arms of the Y-maze contained a water drop. The ants’ decision was noted when they crossed the initial decision line (2cm after arm start) and a final decision line (8cm after arm start, 2cm before end)

### Odour association: Short-term olfactory-memory

#### Experiment 2 – Short-term olfactory-memory: scented runway and flavoured reward versus punishment

The aim of this experiment was to examine if ants can associate odour and flavour cues with a sucrose reward. Flavour here refer to flavouring directly in a food source, while odour refers to environmental odours impregnated into the substrate on which the ant walks to and from a food source. Of course, flavours added to the food source will also have a volatile odour component.

An ant from a donor nest was allowed access to a 10cm-long straight runway covered by a scented paper overlay, which had a drop of matching flavoured sucrose solution at the end (apple or strawberry). While drinking, the ant was marked with a dot of paint on the abdomen, and allowed to return to a recipient nest when satiated. Ants return to recipient nests, not their original donour nest, so as to prevent them from sharing flavoured food with ants which will be tested later. After unloading the sugar, the ant was allowed to return to the apparatus. However, on this visit the linear runway was scented with the other odour, and led to a drop of bitter quinine, also flavoured to match the runway odour. Once the ant had contacted the quinine it was allowed to return to the nest (or gently returned if it did not return on it’s own within 5 minutes). With the training phase concluded, the ant was then allowed onto a Y-maze (as in experiment 1), in which one arm was covered in a reward-associated scented overlay and the other in the punishment-scented overlay. The arm choice of the ant was recorded as in experiment 1 (see “test” in fig. 1b & table 1). 72 ants from three donor colonies were tested, ensuring an equal number of individuals was tested for each odour/flavour and Y-maze side combination daily.

**Table 1:** Experiment and results overview. Experiment 1 involved training over multiple visits on a Y-maze. Experiments 2-9 involved training on a linear runway and testing on a Y-maze with scented arms. Runway odour column and Food Flavour column indicates if the runway was scented or the food flavoured during training, respectivly. In short-term learning experiments training occurred immediately after testing, c. 1-5 minutes. In long-term memory experiments testing occurred the stated number of hours after training. % correct choices are printed in **bold** if significantly different from random choice. In experiments 8 and 9 untrained nestmates, housed with the trained nestmates, were also tested.

| Experiment | Runway odour | Food flavour | Punishment visit (quinine) | N | Short or Long-term memory | % correct choices |
| --- | --- | --- | --- | --- | --- | --- |
| 1) *Side learning: Training on a Y-maze, reward on one arm, punishment on the other (summer)* | - | - | X | 46 | Short | Visit<br>2 = **65%**<br>3 = **84%**<br>4 = **84%**<br>5 = **88%** |
| 2) *scented runway & flavoured reward vs. punishment (summer)* | X | X | X | 72 | Short | **84%** |
| 3) *scented runway & flavoured reward vs. neutral stimulus (summer)* | X | X | - | 48 | Short | **85%** |
| 4) *scented runway & unflavoured reward vs. punishment (summer)* | X | - | X | 72 | Short | **72%** |
| 5) *unscented & flavoured reward vs. neutral stimulus (summer)* | - | X | - | 48 | Short | **62.5%** |
| 6) *unscented & flavoured reward vs. punishment (winter)* | - | X | X | 44 | Short | **86%** |
| 7) *scented runway & unflavoured reward vs. punishment (winter)* | X | - | X | 48 | Short | **87.5%** |
| 8) *scented runway & flavoured reward vs. neutral stimulus (autumn)* | X | X | - | 70 | Long (8, 24, 48 h) | 8h = **85%**<br>24h = **81%**<br>48h = **80%**<br>Nestmates:<br>24h = **77%**<br>48h = **67%** |
| 9) *scented runway & unflavoured reward vs. neutral stimulus (autumn)* | X | - | - | 56 | Long (8, 24, 48 h) | 8h = **67%**<br>24h = **78%**<br>48h = **73%**<br>Nestmates:<br>24h = **40%**<br>48h = **30%** |

#### Experiment 3 – Short-term olfactory-memory: scented runway and flavoured reward versus neutral stimulus

The aim of this experiment was to test whether ants need a positive and a negative stimulus for successful learning, or if only a positive stimulus is sufficient. Ants were trained and tested as in experiment 2 (runway odour and flavoured food), but no second visit to a punished runway was performed. The test was carried out as in experiment 2 (see also fig. 1 & table. 1).

48 ants from two donor colonies were tested, and again both odours/flavours and Y-maze sides were alternated in every combination possible to remove any potential bias.

#### Experiment 4 – Short-term olfactory-memory: scented runway and unflavoured reward versus punishment

The aim of this experiment was to determine whether ants can associate runway odour cues alone with a reward or punishment. Ants were trained and tested as in Experiment 2 (see fig. 1 & table 1), except this time, the food reward and the quinine were not flavoured. 72 ants from 6 donor colonies were tested and both odours and Y-maze sides were alternated daily.

#### Experiment 5 – Short-term olfactory-memory: unscented runway and flavoured reward versus neutral stimulus

The aim of this experiment was to test whether flavoured food, without runway odour or negative reinforcement, is sufficient to form an association between flavoured food and an runway odour presented on a Y-maze arm. This is especially important for potential future applications in pest control. Ants were trained as in experiment 3, except with an unscented runway (see fig. 1 & table.1). In the test, ants had the choice between a scented arm to match the flavour of the flavoured food the ant was trained on, and an unscented one. 48 ants from two donor colonies were tested. Flavours and odour side during tests were balanced.

#### Experiment 6 – Short-term olfactory-memory: unscented runway and flavoured reward versus punishment

The aim of this experiment was to test whether flavoured food, without runway odour, is sufficient to form an association between flavoured food and an runway odour presented on a Y-maze arm, with the addition of punished visit to an alternative odour (unlike in experiment 5, which did not include a punished visit). Ants were trained and tested as in experiment 5 except that a quinine visit was added, as in experiment 2. The test was carried out as in experiment 2 (see fig. 1 & table 1). 44 ants from three donor colonies were tested in winter. Flavours and odour side during tests were balanced.

#### Experiment 7 – Short-term olfactory-memory: scented runway and unflavoured reward versus punishment

The difference between the results of the 5^th^ and 6^th^ experiments were unexpected, considering the results of experiment 3. Moreover, as 5 and 6 were performed in different times of year the conditions were not the same. Here we therefore repeated experiment 4 in the winter to test whether these surprising results were driven by some external factor, possibly related to season or duration of ant captivity (see fig. 1 & table 1). This also allows us to directly compare experiments 6 and 7, both carried out in winter. 48 ants from three donor colonies were tested.

### Odour association: Long-term olfactory-memory

#### Experiment 8 – Long-term olfactory-memory: scented runway and flavoured reward versus neutral stimulus

In this experiment, ants were trained as in experiment 3 (see fig. 1 & table 1) – ants were allowed to make one visit to a flavoured food source via a scented runway. However, every colony was only conditioned to one reward odour. 50 ants were trained per colony. 4 colonies were tested, two of them conditioned to strawberry and two to apple. After training, ants were not allowed back into their original colony or into a recipient colony. Rather, they were housed in a small sub-colony with 60 naïve ants from their donor colony. Trained ants (identifiable as they were marked during training) were tested in a Y-maze, as in experiment 3, after 6, 24, or 48h (4 colonies, 14 ants per period). Each individual ant was only tested once and was then removed. After the trained ants were tested we also tested the untrained nestmates (4 colonies, 20 ants per period, 24h and 48h), to test whether contact with the trained ants, which had fed on flavoured food, allowed them to learn the food flavour and thus follow this odour cue. In total, we tested 70 trained and 80 untrained ants per time period (trained = 6h, 24h and 48h; untrained = 24h and 48h).

#### Experiment 9 – Long-term olfactory-memory: scented runway and unflavoured reward versus neutral stimulus

The aim of this experiment was to exclude the possibility that the ants in experiment 8 were not remembering their association for the whole time period, but rather were refreshing their memory by repeatedly sampling the flavoured food from other nestmates. To this end, this experiment was identical to experiment 8 but with an unflavoured reward; only a scented runway was used (see fig. 1 & table 1). We expected that the trained ants would form an association even without food flavour, but that the untrained ants would not be able to gain any relevant odour information from their sisters. In total, we tested 56 trained and 80 untrained ants from 4 colonies per time period. Two colonies were conditioned to strawberry and two to apple. Y-maze sides containing the conditioned odour were alternated per ant.

### Statistical analysis

Only 3.6% – 10.9% of initial and final choices differed, depending on experiment, so we focused our analysis on the final choices. Data were analysed using generalized linear mixed-effect models (GLMM) (Bolker et al. 2009) in R version 4.1.0 (R Core Team 2021). GLMMs were fitted using the lme4 package (Bates et al. 2015). As the data were binomial (correct / incorrect), a binomial error distribution was used. Since multiple ants were tested per colony, we included colony as random factor. Each model was validated using the DHARMa package (Hartig 2018). Results were plotted using the gglot2 package (Wickham 2016). The complete code and analysis output is provided in supplement 1.

## Results

### Route association: Short-term spatial-memory

#### Experiment 1 – Short-term spatial-memory

This experiment tested whether Argentine ants can associate a side of the Y-maze with the reward (food), and how this memory develops over subsequent visits.

Over all visits, significantly more choices were made for the arm leading to the food (GLMM, n = 46, z-ratio = 2.513 p = 0.045, see fig. 2). In visit 2, after only one visit to the food, 65% (30/46) of ants chose the arm leading to the food (n =46, z-ratio = 2.581, p = 0.042). This rose to 84% (39/46) in visit 3 (n = 46, z-ratio = 1.860, p = 0.026), 84% in visit 4 (38/45) (n = 45, z-ratio = 1.469, p = 0.032) and 88% (39/44) in visit 5 (n = 44, z-ratio = 2.245, p = 0.009). For a pairwise comparison between visits, see online supplement 3, table S1.

**Figure 2:**
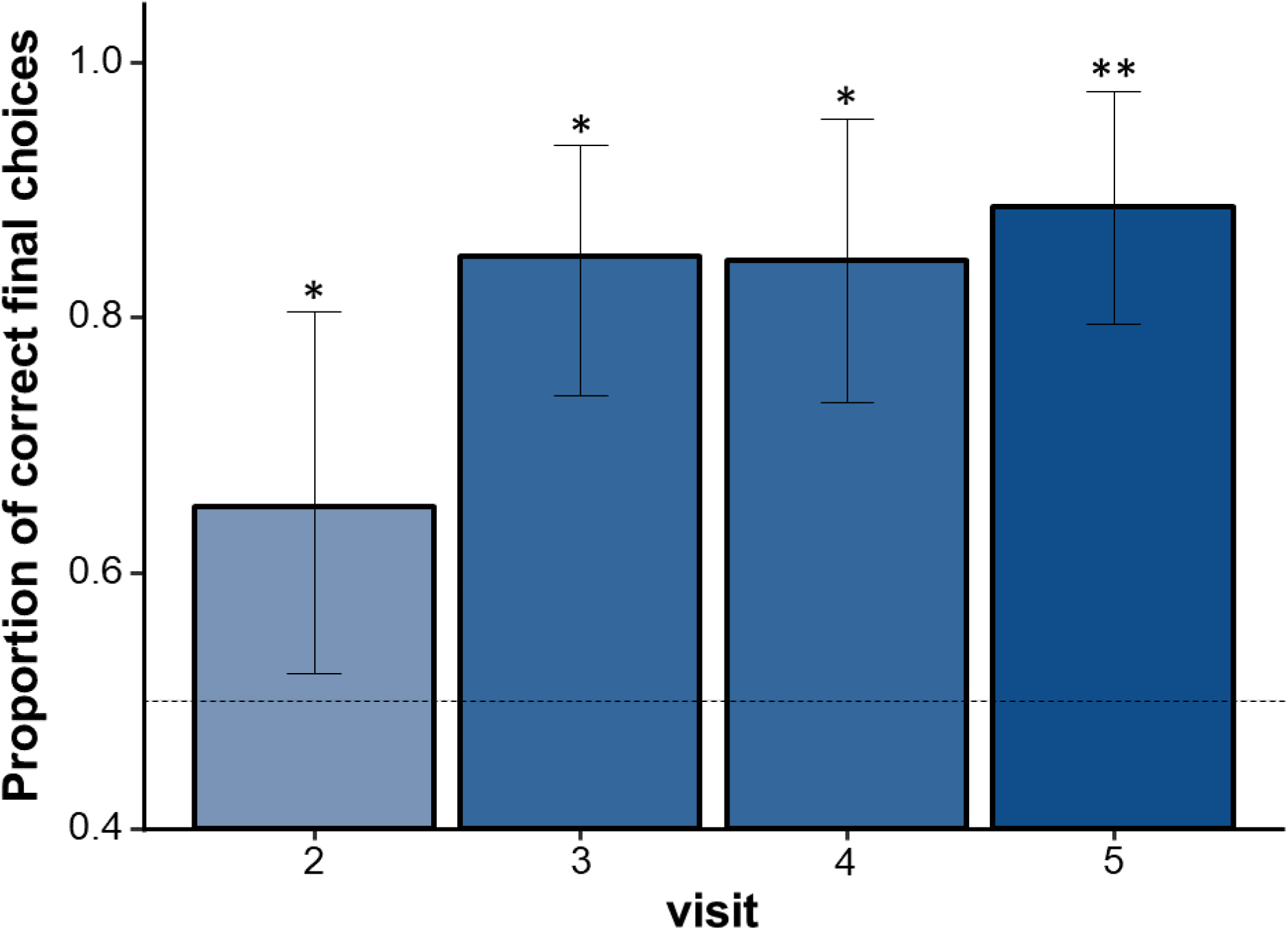
Side-learning: Proportion of ants choosing the side associated with reward (food) per visit. Ants chose the side associated with reward significantly more often than chance after all visits. Bars depict mean, whiskers 95% confidence intervals derived from the fitted GLMM. The dotted horizontal line displays chance level of 50%. \**p* < 0.05; \*\**p* < 0.01; ***p < 0.001.

### Odour association: Short-term olfactory-memory

#### Experiment 2 – 7 Short-term olfactory-memory

In all the experiments there were no differences between the two odours, therefore all results presented are a pooled group of both odours (see supplement 3).

The aim of experiments 2 – 7 was to understand if and how well Argentine ants can associate odours and food flavours with sucrose reward, and whether punishment improves learning (as has been shown in honeybees, (Avarguès-Weber et al. 2010)). The results are summarised in fig. 3 and table 1. In all experiments barring experiment 5, ants chose the correct Y-maze branch significantly more often than would be expected by chance. When comparing experiments 2 and 3, both carried out in the summer, with the only difference between them being the absence of quinine, we see that both show a good learning performance. Thus, there was no evidence that a negative reinforcement (quinine) improved learning. One might be tempted to compare experiments 5 and 6. However, this shoud be avoided, as they were carried out in different conditions. These different conditions, possibly seasonal in nature, can generate differences in behaviour (see differences in experiments 4 and 7).

**Figure 3:**
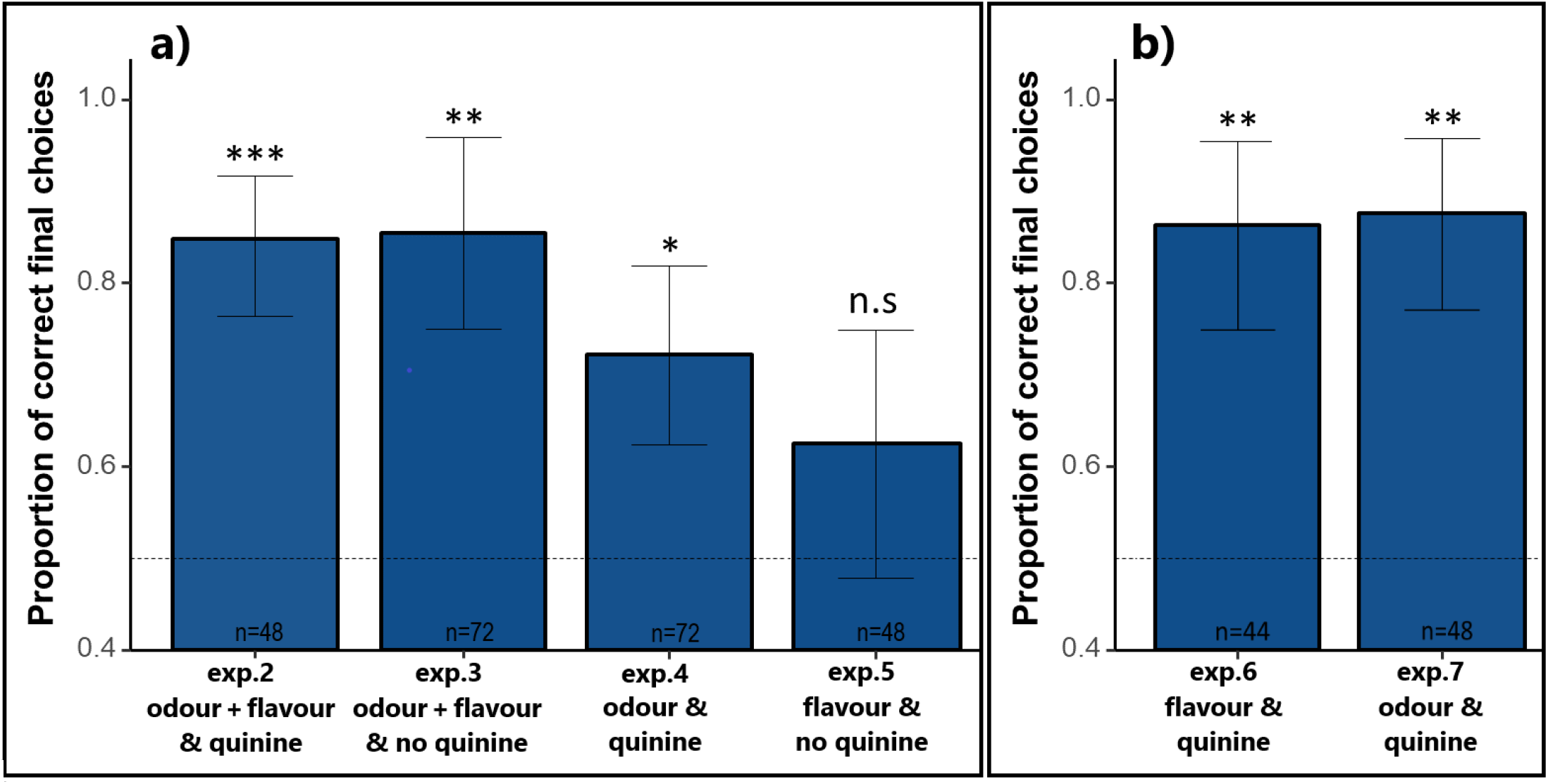
Short-term odour associative learning: Proportion of ants reaching the end of a Y-maze arm scented with an odour associated with a sucrose reward. Ants were able to form a strong association to both training runway odour and food flavour with the reward. Punishment of the contrasting odour using quinine did not improve learning. Experiments shown in different panels were conducted in different seasons (**a** in summer, **b** in winter) and the ants in b had been in captivity longer than in a, and therefore should not be compared. Bars depict means, whiskers 95% confidence intervals derived from the fitted GLMM. The dotted horizontal line displays chance level of 50%. \**p* < 0.05; \*\**p* < 0.01; ***p < 0.001.

There is evidence that combining path odour and food flavour (experiment 2) improved learning compared to using only the odour of the runway (experiment 4). The same pattern is evident in that the combination of path odour and food flavor (exp. 5) was better than only flavour (exp. 3). Therefore, using two cues leads to better performance than using just one. For a detailed description of these results, see supplement 4.

Experiment 6 and 7 showed that only odour on the runway or only flavour in the food can drive for very similar and good learning performances. Thus, only one cue is enough for establishing short term memories.

### Odour-association: Long-term olfactory-memory

#### Experiment 8 – 9 Long-term olfactory-memory

The aim of experiments 8 and 9 was to examine if Argentine ants also form long-term memories (6h, 24h, 48h) associating an odour (of the runway and of the food) with the reward (experiment 8); or only the odour of the runway with a reward (experiment 9). The results are summarised in fig. 4 and table 1. Ants chose the correct branch even after 48h. Untrained ants exposed to flavoured food via trained ants also chose the correct branch after 24 and 48h, but untrained ants housed with ants trained on unflavoured food showed no preference for the correct odour after 24h, and avoided it after 48h. Detailed results are provided in supplement 4.

**Figure 4:**
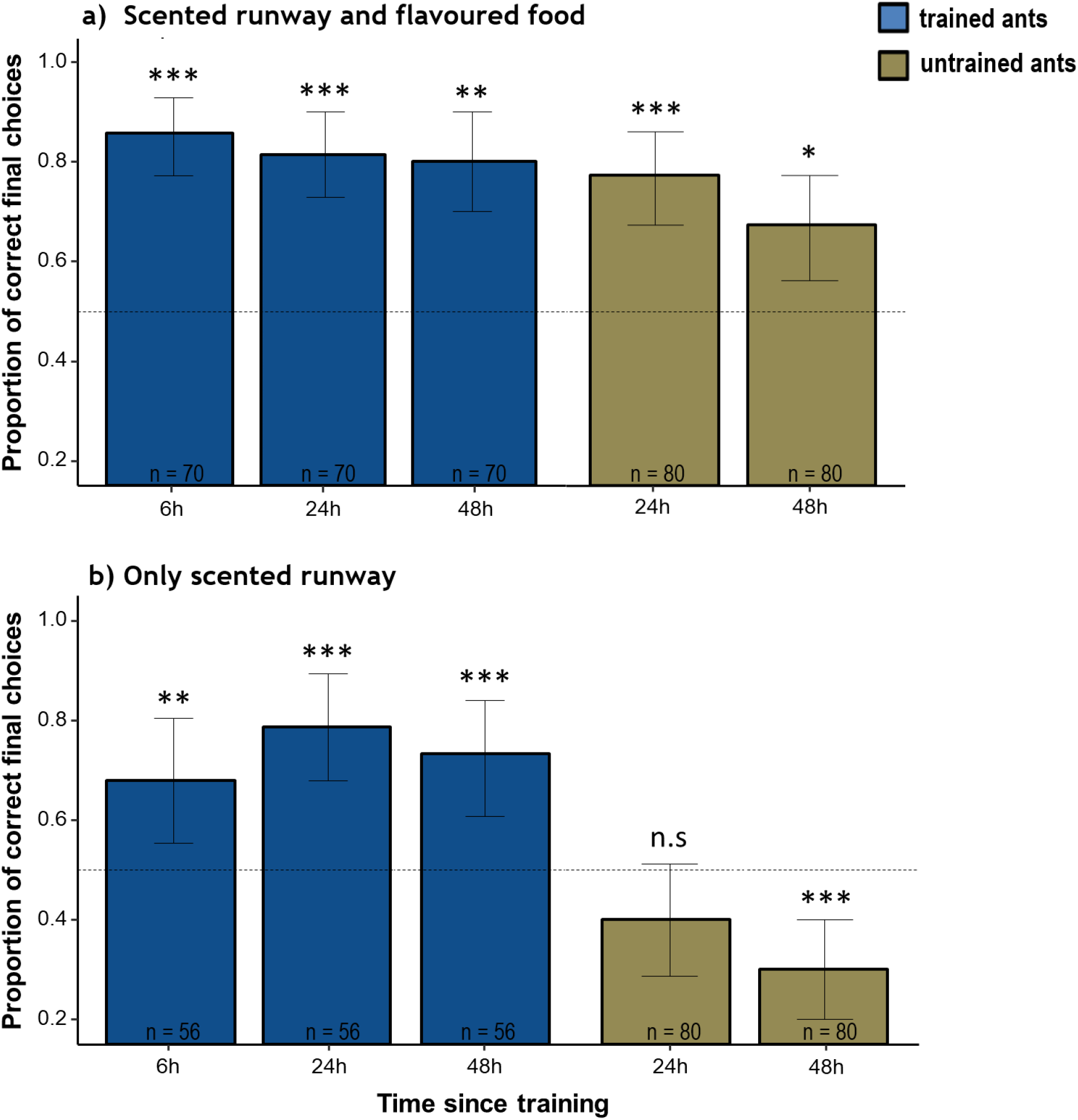
Long-term odour associative learning: Proportion of ants choosing the odour associated with a sucrose reward. Trained ants were tested after 6h, 24h, and 48h. **(a):** When the runway was scented and food flavoured, the trained ants learned the association and preferred the odour previously associated with sucrose. Untrained nestmates also showed a preference for the rewarded odour when food was flavoured, which was not significantly weaker than that of trained ants. **(b):** When only the runway was scented during training, and food was unflavoured, trained ants were able to establish an association and then choose the correct odour after 6h, 24h and 48h. By contrast, untrained nestmates did not show any odour preference after 24h, and showed an aversion for the food odour after 48h. Bars depict means, whiskers 95% confidence intervals derived from the fitted GLMM. The dotted horizontal line displays chance level of 50%. \**p* < 0.05; \*\**p* < 0.01; ***p < 0.001.

## Discussion

*Linepithema humile* are adept learners: The side-learning results show that Argentine ants are capable of learning to associate a direction or an arm of a Y-maze with a food reward after just one visit, and accuracy improves with further visits (see fig. 2). This finding has also recently been replicated in a separate study (Galante and Czaczkes 2022). Such rapid and reliable learning is surprising, given the fact that *L. humile* relies strongly on social information (pheromone trails), while other recruiting social insects which learn equally fast, such as the ants *Lasius niger* and *Paraponera clavata* ants, and the honeybee *Apis melifera*, prioritize memories over recruitment signals when the two conflict (Harrison et al. 1989; Aron et al. 1993; Grüter et al. 2008, 2011; Czaczkes et al. 2013; von Thienen et al. 2016). *L. humile’s* strong individual memory may allow them to reliably recruit even to distant resources, while their strong reliance on social information (von Thienen et al. 2014, 2016), and ability to recruit from active trails (Flanagan et al. 2013), may allow them to rapidly dominate these resources once found. Indeed, the rapid discovery of new resources coupled with massive, rapid recruitment might be an important contribution to making Argentine ants competitive against native ants (Holway 1999), but see (Cordonnier et al. 2020). Route learning and navigation in ants is less effective on complex routes with alternating turns, but when trail pheromones are present, error rates decrease (Czaczkes et al. 2013). The combination of social information and a strong individual side-learning memory could be beneficial and explain why Argentine ants developed such a strong individual route learning ability in parallel to their pheromone recruitment system. However, a systematic exploration of learning in (social) insects has not been carried out, so it is also possible that many or most ants are equally good learners.

Alongside their impressive route learning, Argentine ants show even more rapid learning of olfactory cues. Ants were able to form a strong association after only one rewarded visit (see fig. 3 & table 1). These findings are again comparable to results from *La. niger* (Czaczkes et al. 2014; Czaczkes and Kumar 2020), and were also recently replicated in a separate study on *L. humile* (Galante and Czaczkes 2022). Somewhat surprisingly, results from both the short-term and long-term learning experiments showed that punishing the alternative odour with quinine does not seem to have an strong impact on learning. The lack of effect of punishment is further supported by the lack of difference in learning between experiments 2 (punished) and 3 (unpunished). Finally, the excellent learning found in the long-term learning experiments (only odour, no quinine) again suggest that punishment is not nececcary for strong learning, and that the results of experiment 5 (only flavour, no quinine) are an underestimate. Indeed, *Camponotus fellah* ants failed to form a negative association with an alternative odour in a discrimination test (Josens et al. 2009), although studies on *La. niger* ants and honeybees show rapid association of odours with negative reinforcement (Avarguès-Weber et al. 2010; Wenig et al. 2021). Caution is required when comparing the apparently large improvement in learning in experiment 6 (flavour & quinine) versus experiment 5 (flavour, no quinine) since, as started previously, those experiments were performed in different seasons, with ants which had spend different amounts of time in captivity, and therefore in different conditions. The differences observed suggested that some difference between these experimental periods resulted in better performances. We explored this by repeating experiment 4 in the winter (experiment 7), and indeed found again much improved learning (see supplement 4, experiment 4 and 7). It might be that these ants learn better in winter. While this would be surprising, *L. humile* behaviour has been shown to vary seasonally (Sanmartín-Villar et al. 2022). It is important to note, however, that this study was not designed to test any seasonal effect. Performance differences between experiments performed at different times may be due to season, time in captivity, or some other factors.

Especially remarkable is that *L. humile* is able to form an association with a sucrose reward given only one cue: a food flavour (experiment 6) or a runway odour (experiments 7 & 9). This is convenient if associative learning is to be applied to improving *L. humile* control in the field, where flavouring baits is straightforward, but adding enviromental odours not. While learning of single cues can be very strong, offering multiple cues could have an additive or sub-additive effect on learning, especially under conditions where learning is pronounced (see figure 3a).

To our knowledge, the learning ability of *Linepithema humile* has never before been described in such detail, and their very rapid learning of olfactory cues never fully appreciated. Rossi et al. (2020) previously demonstrated that *L. humile* could associate an odour with a food reward. Our results support their findings using a different experimental approach, and went further in exploring the cognitive abilities of this species with a higher resolution: the Rossi study allowed 3 visits to a circular open foraging arena with only one odour presented with the positive reward, while our discrimination paradigm (odour 1 vs. odour 2) was more demanding for the ants. Furthermore, our study controlled the experiences of the individual ants more tightly, by reducing the number of rewarded visits to one, and separating rewarded and unrewarded learning events. To our knowledge, our study is also the first study to demonstrated that *L. humile* can learn to associate the flavour of the food alone with the food reward.

Alongside the strong short-term olfactory memory, Argentine ants also possess a strong long-term memory, which lasts for at least 48h (see fig. 4 & table 1). As in the short-term memory tests, a runway odour alone, without food flavour, was sufficient to drive strong and stable learning. This is in line with studies on other species (*Formica fusca* and *Camponotus fellah)*, which show stable memory for at least 72h, and a decay in learning after a week (Josens et al. 2009; Piqueret et al. 2019). However, the study on *Camponotus fellah* used very extensive training (16 training visits). We found that one exposure alone, without a punishment visit, was sufficient to elicit strong and stable learning.

In the long-term memory experiments, tested ants were housed in a small sub-nest where they fed their nestmates after their training. As a result, flavoured food was likely distributed amongst the workers via trophallaxis. Inclusion of food flavour resulted in the untrained nestmates also developing a preference for that odour, demonstrating a very effective distribution of food-related information within the nest. Such intranidal information spread affecting future foraging decision has been reported in both other ants (Roces 1990; Provecho and Josens 2009; Arenas and Roces 2018), bees (Farina et al. 2005) and wasps (Jeanne and Taylor 2009).

*Linepithema humile* are adept learners and rapidly disseminate food-related information in the nest – abilities which might help them to adapt to new environments and dominate resources, and could be one reason for their success as invaders. However, their strength could potentially become a weakness, if their memory can be used against them. One possibility would be to use associative learning and intranidal information transfer to steer foraging preference towards poisoned baits (Josens et al. 2016), much as honeybee preference can be steered for pollination purposes (Farina et al. 2020b). While leveraging learning to control animal behaviour is commonly used in behavioural conservation of vertebrates, it is mostly used to reduce crop damage or to minimize negative interactions with humans (Matsuzawa et al. 1983; Webb et al. 2015; Valenta et al. 2021). Leveraging learning to improve the control of invasive animals may be an underappreciated tool in the conservation toolbox.

## Supporting information

Supplements 1 - R Markdown

Supplements 2 - Complete raw data file

Supplements 3 - Pilot-Study and Side-learning pairwise tabletrash

Supplements 4 - Detailed results with statistics AB

## Acknowledgements

We would like to thank Silvia Abril and Eduardo Sequeira for providing ant colonies. Thomas Wagner and Henrique Galante were supported by an ERC starter grant (Cognitive control: 948181) to Tomer J. Czaczkes. Tomer J. Czaczkes was supported by a Heisenberg fellowship from the Deutsche Forschungsgemeinschaft (CZ237/4-1). R.J thanks the CONICET of Argentina, and the ANPCyT (PICT 2017-9) for funding.

## Author contribution

T. Wagner, H. Galante and T. J. Czaczkes designed the study. T. Wagner and H. Galante collected the data. T. Wagner analysed the data and wrote the manuscript. H. Galante, R. Josens, and T. J. Czaczkes edited the manuscript. All authors read and approved the manuscript.

