## Supplements 1 - R Markdown for "A systematic examination of learning in the invasive ant *Linepithema humile* reveals very rapid development of short and long-term memory"

#### 15 10 2022

### ESM 1 - Introduction

This supplement provides the entire R script and output of the
statistical analysis we performed and figures produced, in their
original form. It is presented in the spirit of open and transparent
science, but has not been carefully curated.

Here, we conduct a comprehensive in-depth investigation of learning
in the ecologically important ant L. humile. We study how rapidly L.
humile form short- and long-term memories, which types of cues best
support this, and whether food-related cues are transferred between
nestmates intranidally. We split the experiments into 3 different parts.
1: Side-learning 2: Odour associative learning “short term”. 3: Odour
associative learning “long term”

### Data import, libraries, housekeeping

loading libraries

```
library (ggplot2)
library (gplots)
library (ggpubr)
library (lme4)
library (nlme) 
library (tidyverse)
library (emmeans)
library (multcomp)
library (DHARMa)
library (readxl)
library (wesanderson)
library (grid)
library (gridExtra)
library (dplyr)
library (scales)
library (lattice)
library (MASS)
library(EnvStats)
library(officer)
library(rvg)
library(Hmisc)
library(eoffice)
library(cowplot)
```

setting directory

import all data files for later analysis. For privacy echo=
FALSE.

used datasets:

PreferenceOdour1vsOdour2

Exp1SideLearning

Exp2FlavourOdourQuinine

Exp3FlavourOdourNoQuinine

exp2\_exp3\_merged

Exp4OnlyOdourQuinine

Exp5OnlyFlavourNoQuinine

Exp6OnlyFlavourQuinineWinter

exp5\_exp6\_merged

Exp7OnlyOdourQuinineWinter

exp4\_exp7\_merged

Exp8LongTermFlavourOdourNoQuinineTrained

Exp8LongTermFlavourOdourNoQuinineNestmates

Exp8LongTermFlavourOdourNoQuinineTrainedVsNestmates

Exp9LongTermOnlyOdourNoQuinineTrained

Exp9LongTermOnlyOdourNoQuinineNestmates

Exp9LongTermFlavourOdourNoQuinineTrainedVsNestmates

### Statistical analysis

#### 0. Analysis, pilot study odour preference test

Ants were tested on a Y-maze for their odour preference. Both Y-maze
arms were covered in scented paper overlays (e.g left strawberry and
right apple). Data is binomial distributed.

n = 158 tested ants, 92 chose strawberry, binomial choice. Result:
Ants chose significantly more often strawberry over apple. Nevertheless,
the difference seems to be minor and through our alternately odour
treatment 50% reward odour = apple, and vice versa were decided to use
these two odours.

##### Fast binomial test, pilot study

```
binom.test(92,158,0.5)
```

```
## 
##  Exact binomial test
## 
## data:  92 and 158
## number of successes = 92, number of trials = 158, p-value = 0.04637
## alternative hypothesis: true probability of success is not equal to 0.5
## 95 percent confidence interval:
##  0.5012756 0.6601356
## sample estimates:
## probability of success 
##              0.5822785
```

#### 1. Analysis, side-learning experiment

Can L. humile ants learn to associate a location, one arm of a
Y-maze, with the presence of a reward? Here we investigate this by
offering a reward, sucrose, on one of the Y-maze arms and a punishment,
quinine, on the opposing arm. Data is binomial distributed and colony
ID, ant ID as random factor.

##### Glmm modeling & pairwise, experiment 1

```
# Glmm

Exp1SideLearning$visit<-as.factor(Exp1SideLearning$visit)

Exp1SideLearning_m <-glmer(correct_final ~ visit + (1|colony/antid),
  family="binomial",
  data= Exp1SideLearning)

summary(Exp1SideLearning_m)
```

```
## Generalized linear mixed model fit by maximum likelihood (Laplace
##   Approximation) [glmerMod]
##  Family: binomial  ( logit )
## Formula: correct_final ~ visit + (1 | colony/antid)
##    Data: Exp1SideLearning
## 
##      AIC      BIC   logLik deviance df.resid 
##    178.1    197.3    -83.0    166.1      175 
## 
## Scaled residuals: 
##     Min      1Q  Median      3Q     Max 
## -2.9096  0.2533  0.3081  0.4180  1.0007 
## 
## Random effects:
##  Groups       Name        Variance Std.Dev.
##  antid:colony (Intercept) 0.8643   0.9297  
##  colony       (Intercept) 0.0000   0.0000  
## Number of obs: 181, groups:  antid:colony, 46; colony, 6
## 
## Fixed effects:
##             Estimate Std. Error z value Pr(>|z|)   
## (Intercept)   0.7544     0.3777   1.997  0.04579 * 
## visit3        1.2519     0.5630   2.223  0.02618 * 
## visit4        1.1958     0.5591   2.139  0.03244 * 
## visit5        1.5871     0.6131   2.589  0.00963 **
## ---
## Signif. codes:  0 '***' 0.001 '**' 0.01 '*' 0.05 '.' 0.1 ' ' 1
## 
## Correlation of Fixed Effects:
##        (Intr) visit3 visit4
## visit3 -0.480              
## visit4 -0.494  0.405       
## visit5 -0.440  0.379  0.375
## optimizer (Nelder_Mead) convergence code: 0 (OK)
## boundary (singular) fit: see help('isSingular')
```

```
# Pairwise

Pairwise1 = emmeans(Exp1SideLearning_m, spec= "visit")

Pairwise1=contrast(Pairwise1, method = "pairwise")
summary(Pairwise1)
```

```
##  contrast        estimate    SE  df z.ratio p.value
##  visit2 - visit3  -1.2519 0.563 Inf  -2.223  0.1169
##  visit2 - visit4  -1.1958 0.559 Inf  -2.139  0.1408
##  visit2 - visit5  -1.5871 0.613 Inf  -2.589  0.0474
##  visit3 - visit4   0.0561 0.612 Inf   0.092  0.9997
##  visit3 - visit5  -0.3353 0.657 Inf  -0.511  0.9566
##  visit4 - visit5  -0.3913 0.657 Inf  -0.596  0.9334
## 
## Results are given on the log odds ratio (not the response) scale. 
## P value adjustment: tukey method for comparing a family of 4 estimates
```

Does this model work?

```
dharmit1 <-simulateResiduals(Exp1SideLearning_m)
plot(dharmit1)
```

Model works.

##### Figures

```
#final decision#

fig1 <- ggplot (Exp1SideLearning, aes(x = visit, y = correct_final))+
  scale_y_continuous(expand = c(0, 0)) + # forces X axis to 0, but in this case is overriden by ribbon
  geom_point( alpha = 0) +
  stat_summary(fun.y = "mean", geom = "bar", fill="dodgerblue4") +
  ylab("Proportion of correct final choices") +
  xlab("visit") + stat_summary(fun.data = "mean_cl_boot", geom="errorbar", width = 0.2) +
  theme_bw(18) +
  geom_abline(intercept = 0.5, slope = 0, color = "black", linetype = 2) +
  coord_cartesian( ylim = c(0.4, 1.05)) # zooms in on the top half

print(fig1)
```

Ants learned already after the first real test visit (visit 2) to
associate a side with the reward. Only the 5th visit was significantly
stronger than the 2nd visit.

### Odour-flavour-learning shot-term

#### 2. Analysis odour and flavour and quinine (experimental set 1, summer)

Can ants learn correctly to associate the reward odour and flavour
with the reward? Reward visit (sucrose) and punishment visit (quinine).
Reward odour is apple 50% or strawberry 50%, Y- maze reward odour 1
vs. odour 2. Data is binomial distributed and colony ID as random
factor.

##### Glmm modeling & pairwise, experiment 2

```
# Glmm

Exp2FlavourOdourQuinine$Reward_Odour<-as.factor(Exp2FlavourOdourQuinine$Reward_Odour)


Exp2FlavourOdourQuinine_m <-glmer(correct_final ~ Reward_Odour + (1|Colony_ID_donor),
                              family="binomial",
                              data= Exp2FlavourOdourQuinine)
```

```
## boundary (singular) fit: see help('isSingular')
```

```
summary(Exp2FlavourOdourQuinine_m)
```

```
## Generalized linear mixed model fit by maximum likelihood (Laplace
##   Approximation) [glmerMod]
##  Family: binomial  ( logit )
## Formula: correct_final ~ Reward_Odour + (1 | Colony_ID_donor)
##    Data: Exp2FlavourOdourQuinine
## 
##      AIC      BIC   logLik deviance df.resid 
##     66.6     73.4    -30.3     60.6       69 
## 
## Scaled residuals: 
##     Min      1Q  Median      3Q     Max 
## -2.8284  0.3535  0.3535  0.4913  0.4913 
## 
## Random effects:
##  Groups          Name        Variance Std.Dev.
##  Colony_ID_donor (Intercept) 0        0       
## Number of obs: 72, groups:  Colony_ID_donor, 3
## 
## Fixed effects:
##                        Estimate Std. Error z value Pr(>|z|)    
## (Intercept)              1.4214     0.4211   3.375 0.000737 ***
## Reward_OdourStrawberry   0.6581     0.6772   0.972 0.331179    
## ---
## Signif. codes:  0 '***' 0.001 '**' 0.01 '*' 0.05 '.' 0.1 ' ' 1
## 
## Correlation of Fixed Effects:
##             (Intr)
## Rwrd_OdrStr -0.622
## optimizer (Nelder_Mead) convergence code: 0 (OK)
## boundary (singular) fit: see help('isSingular')
```

```
# Pairwise 

Pairwise2 = emmeans(Exp2FlavourOdourQuinine_m, spec= "Reward_Odour")

Pairwise2=contrast(Pairwise2, method = "pairwise")
summary(Pairwise2)
```

```
##  contrast           estimate    SE  df z.ratio p.value
##  Apple - Strawberry   -0.658 0.677 Inf  -0.972  0.3312
## 
## Results are given on the log odds ratio (not the response) scale.
```

Does the model work?

```
dharmit2 <-simulateResiduals(Exp2FlavourOdourQuinine_m)
plot(dharmit2)
```

Ants learned to associate two cues (flavour and odour) with the food
reward in a reward-punishment setup.

#### 3. Analysis odour and flavour no quinine (experimental set 1, summer)

Can ants learn correctly to associate the reward odour and flavour
with the reward, without having a punishment visit (quinine). Reward
odour is apple 50% or strawberry 50%, Y- maze reward odour 1 vs. odour
2. Data is binomial distributed and colony ID as random factor.

##### Glmm modeling & pairwise, experiment 3

```
# Glmm

Exp3FlavourOdourNoQuinine$Reward_Odour<-as.factor(Exp3FlavourOdourNoQuinine$Reward_Odour)


Exp3FlavourOdourNoQuinine_m <-glmer(correct_final ~ Reward_Odour + (1|Colony_ID_donor),
                              family="binomial",
                              data= Exp3FlavourOdourNoQuinine)
```

```
## boundary (singular) fit: see help('isSingular')
```

```
summary(Exp3FlavourOdourNoQuinine_m)
```

```
## Generalized linear mixed model fit by maximum likelihood (Laplace
##   Approximation) [glmerMod]
##  Family: binomial  ( logit )
## Formula: correct_final ~ Reward_Odour + (1 | Colony_ID_donor)
##    Data: Exp3FlavourOdourNoQuinine
## 
##      AIC      BIC   logLik deviance df.resid 
##     45.7     51.3    -19.9     39.7       45 
## 
## Scaled residuals: 
##     Min      1Q  Median      3Q     Max 
## -2.6458  0.3780  0.3780  0.4472  0.4472 
## 
## Random effects:
##  Groups          Name        Variance Std.Dev.
##  Colony_ID_donor (Intercept) 0        0       
## Number of obs: 48, groups:  Colony_ID_donor, 2
## 
## Fixed effects:
##                        Estimate Std. Error z value Pr(>|z|)   
## (Intercept)              1.6094     0.5477   2.938   0.0033 **
## Reward_OdourStrawberry   0.3365     0.8252   0.408   0.6835   
## ---
## Signif. codes:  0 '***' 0.001 '**' 0.01 '*' 0.05 '.' 0.1 ' ' 1
## 
## Correlation of Fixed Effects:
##             (Intr)
## Rwrd_OdrStr -0.664
## optimizer (Nelder_Mead) convergence code: 0 (OK)
## boundary (singular) fit: see help('isSingular')
```

```
# Pairwise

Pairwise3 = emmeans(Exp3FlavourOdourNoQuinine_m, spec= "Reward_Odour")

Pairwise3=contrast(Pairwise3, method = "pairwise")
summary(Pairwise3)
```

```
##  contrast           estimate    SE  df z.ratio p.value
##  Apple - Strawberry   -0.336 0.825 Inf  -0.408  0.6835
## 
## Results are given on the log odds ratio (not the response) scale.
```

Figures of experiment 2 & 3 were merged for a better comparison
in this script. For the manuscript,all short-term experiments were
merged to one big figure. Aesthetic judgement were made via
powerpoint.

##### Figures merged Experiment 2 & 3

```
# Base plot

exp2_exp3_merged_final_plot <- ggplot (exp2_exp3_merged, aes(x = Reward_Odour, y = correct_final))+
  scale_y_continuous(expand = c(0, 0)) + # forces X axis to 0, but in this case is overriden by ribbon
  geom_point( alpha = 0) +
  stat_summary(fun.y = "mean", geom = "bar", fill="dodgerblue4") +
  stat_summary(fun.data = "mean_cl_boot", geom="errorbar", width = 0.2) +
  ylab("Proportion of correct final choices") +
  xlab("threatment") +
  theme_bw(26) + theme(axis.line = element_line(colour = "black"),
    panel.grid.major = element_blank(),
    panel.grid.minor = element_blank(),
    panel.border = element_blank(),
    panel.background = element_blank()) +
  geom_abline(intercept = 0.5, slope = 0, color = "black", linetype = 2) +
  coord_cartesian( ylim = c(0.4, 1.05)) # zooms in on the top half
```

```
## Warning: `fun.y` is deprecated. Use `fun` instead.
```

```
print(exp2_exp3_merged_final_plot)
```

```
# Powerpoint import

fig_exp2_exp3_final_print <- "fig_exp2_exp3_print.pptx"

fig_exp2_exp3_final_pp <- dml(ggobj = exp2_exp3_merged_final_plot)

read_pptx()%>%
add_slide() %>%
ph_with(fig_exp2_exp3_final_pp, location = ph_location_fullsize())%>%
print(fig_exp2_exp3_final_print)

ggsave ("exp2_exp3_merged_final.png", plot = exp2_exp3_merged_final_plot, dpi = 300, width = 20, height = 20, units = c("cm"))
```

Ants were able, even without having a punishment visit, to associate
two cues (flavour and odour) with the food reward.

#### 4. Analysis, odour no flavour and quinine (summer)

Can ants learn correctly to associate the reward odour alone, without
having food flavour but with having a punishment visit (quinine), with
the reward. Reward odour is apple 50% or strawberry 50%, Y- maze reward
odour 1 vs. odour 2. Data is binomial distributed and colony ID as
random factor.

##### Glmm modeling & pairwise, experiment 4

```
# Glmm

Exp4OnlyOdourQuinine$Reward_Odour<-as.factor(Exp4OnlyOdourQuinine$Reward_Odour)


Exp4OnlyOdourQuinine_m <-glmer(correct_final ~ Reward_Odour + (1|Colony_ID),
                                family="binomial",
                                data= Exp4OnlyOdourQuinine)
```

```
## boundary (singular) fit: see help('isSingular')
```

```
summary(Exp4OnlyOdourQuinine_m)
```

```
## Generalized linear mixed model fit by maximum likelihood (Laplace
##   Approximation) [glmerMod]
##  Family: binomial  ( logit )
## Formula: correct_final ~ Reward_Odour + (1 | Colony_ID)
##    Data: Exp4OnlyOdourQuinine
## 
##      AIC      BIC   logLik deviance df.resid 
##     91.1     97.9    -42.5     85.1       69 
## 
## Scaled residuals: 
##     Min      1Q  Median      3Q     Max 
## -1.6124 -1.6124  0.6202  0.6202  0.6202 
## 
## Random effects:
##  Groups    Name        Variance Std.Dev.
##  Colony_ID (Intercept) 0        0       
## Number of obs: 72, groups:  Colony_ID, 6
## 
## Fixed effects:
##                         Estimate Std. Error z value Pr(>|z|)  
## (Intercept)            9.555e-01  3.721e-01   2.568   0.0102 *
## Reward_OdourStrawberry 4.919e-16  5.262e-01   0.000   1.0000  
## ---
## Signif. codes:  0 '***' 0.001 '**' 0.01 '*' 0.05 '.' 0.1 ' ' 1
## 
## Correlation of Fixed Effects:
##             (Intr)
## Rwrd_OdrStr -0.707
## optimizer (Nelder_Mead) convergence code: 0 (OK)
## boundary (singular) fit: see help('isSingular')
```

```
# Pairwise

Pairwise4 = emmeans(Exp4OnlyOdourQuinine_m, spec= "Reward_Odour")

Pairwise4=contrast(Pairwise4, method = "pairwise")
summary(Pairwise4)
```

```
##  contrast            estimate    SE  df z.ratio p.value
##  Apple - Strawberry -4.92e-16 0.526 Inf   0.000  1.0000
## 
## Results are given on the log odds ratio (not the response) scale.
```

#### 7. Analysis, Odour no flavour and quinine (winter)

We received additional data, this time in winter. Again, only odour
and quinine, but no flavour. Reward odour is apple 50% or strawberry
50%, Y- maze reward odour 1 vs. odour 2. Data is binomial distributed
and colony ID as random factor.

##### Glmm modeling & pairwise, experiment 7

we called exp 4b exp 7 in the working process (file names in R)
because we wanted to test for a season effect but so it was excluded as
another exp. After feeding we merged the data

```
# Glmm

Exp7OnlyOdourQuinineWinter$Reward_Odour<-as.factor(Exp7OnlyOdourQuinineWinter$Reward_Odour)

Exp7OnlyOdourQuinineWinter_m <-glmer(correct_final ~ Reward_Odour + (1|Colony_ID),
                                  family="binomial",
                                  data= Exp7OnlyOdourQuinineWinter)
```

```
## boundary (singular) fit: see help('isSingular')
```

```
summary(Exp7OnlyOdourQuinineWinter_m)
```

```
## Generalized linear mixed model fit by maximum likelihood (Laplace
##   Approximation) [glmerMod]
##  Family: binomial  ( logit )
## Formula: correct_final ~ Reward_Odour + (1 | Colony_ID)
##    Data: Exp7OnlyOdourQuinineWinter
## 
##      AIC      BIC   logLik deviance df.resid 
##     38.9     44.5    -16.4     32.9       45 
## 
## Scaled residuals: 
##     Min      1Q  Median      3Q     Max 
## -4.7958  0.2085  0.2085  0.5130  0.5130 
## 
## Random effects:
##  Groups    Name        Variance Std.Dev.
##  Colony_ID (Intercept) 0        0       
## Number of obs: 48, groups:  Colony_ID, 3
## 
## Fixed effects:
##                        Estimate Std. Error z value Pr(>|z|)   
## (Intercept)              1.3350     0.5026   2.656  0.00791 **
## Reward_OdourStrawberry   1.8005     1.1385   1.582  0.11376   
## ---
## Signif. codes:  0 '***' 0.001 '**' 0.01 '*' 0.05 '.' 0.1 ' ' 1
## 
## Correlation of Fixed Effects:
##             (Intr)
## Rwrd_OdrStr -0.441
## optimizer (Nelder_Mead) convergence code: 0 (OK)
## boundary (singular) fit: see help('isSingular')
```

```
# Pairwise

Pairwise7 = emmeans(Exp7OnlyOdourQuinineWinter_m, spec= "Reward_Odour")

Pairwise7=contrast(Pairwise7, method = "pairwise")
summary(Pairwise7)
```

```
##  contrast           estimate   SE  df z.ratio p.value
##  Apple - Strawberry     -1.8 1.14 Inf  -1.582  0.1138
## 
## Results are given on the log odds ratio (not the response) scale.
```

Figures of experiment 4 & 7 were merged for a better comparison
in this script. For the manuscript,all short-term experiments were
merged to one big figure. Aesthetic judgement were made via
powerpoint.

##### Figures merged Experiment 4 & 7

```
# Basic plot

exp4_exp7_merged_final_plot <- ggplot (exp4_exp7_merged, aes(x = Reward_Odour, y = correct_final))+
  scale_y_continuous(expand = c(0, 0)) + # forces X axis to 0, but in this case is overriden by ribbon
  geom_point( alpha = 0) +
  stat_summary(fun.y = "mean", geom = "bar", fill="dodgerblue4") +
  stat_summary(fun.data = "mean_cl_boot", geom="errorbar", width = 0.2) +
  ylab("correct choices") +
  xlab("threatment") +
  theme_bw(26) + theme(axis.line = element_line(colour = "black"),
    panel.grid.major = element_blank(),
    panel.grid.minor = element_blank(),
    panel.border = element_blank(),
    panel.background = element_blank())  +
  geom_abline(intercept = 0.5, slope = 0, color = "black", linetype = 2) +
  coord_cartesian( ylim = c(0.4, 1.05)) # zooms in on the top half
print(exp4_exp7_merged_final_plot)
```

```
# Powerpoint import


fig_exp4_exp7_final_print <- "fig_exp4_exp7_print.pptx"

fig_exp4_exp7_final_pp <- dml(ggobj = exp4_exp7_merged_final_plot)

read_pptx()%>%
add_slide() %>%
ph_with(fig_exp4_exp7_final_pp, location = ph_location_fullsize())%>%
print(fig_exp4_exp7_final_print)

ggsave ("exp4_exp7_merged_final.png", plot = exp4_exp7_merged_final_plot, dpi = 300, width = 20, height = 20, units = c("cm"))
```

##### Glmm modeling & pairwise comparison experiment 4 vs 7

```
# Glmm

exp4_exp7_final_model <-glmer(correct_final ~ Reward_Odour + (1|Colony_ID),
                                  family="binomial",
                                  data= exp4_exp7_merged)
```

```
## boundary (singular) fit: see help('isSingular')
```

```
summary(exp4_exp7_final_model)
```

```
## Generalized linear mixed model fit by maximum likelihood (Laplace
##   Approximation) [glmerMod]
##  Family: binomial  ( logit )
## Formula: correct_final ~ Reward_Odour + (1 | Colony_ID)
##    Data: exp4_exp7_merged
## 
##      AIC      BIC   logLik deviance df.resid 
##    127.3    135.6    -60.6    121.3      117 
## 
## Scaled residuals: 
##     Min      1Q  Median      3Q     Max 
## -2.6458  0.3780  0.3780  0.6202  0.6202 
## 
## Random effects:
##  Groups    Name        Variance Std.Dev.
##  Colony_ID (Intercept) 0        0       
## Number of obs: 120, groups:  Colony_ID, 9
## 
## Fixed effects:
##                          Estimate Std. Error z value Pr(>|z|)    
## (Intercept)                0.9555     0.2631   3.632 0.000282 ***
## Reward_OdourOdourQWinter   0.9904     0.5096   1.943 0.051963 .  
## ---
## Signif. codes:  0 '***' 0.001 '**' 0.01 '*' 0.05 '.' 0.1 ' ' 1
## 
## Correlation of Fixed Effects:
##             (Intr)
## Rwrd_OdrOQW -0.516
## optimizer (Nelder_Mead) convergence code: 0 (OK)
## boundary (singular) fit: see help('isSingular')
```

```
# Pairwise

Pairwise_exp4_exp7 = emmeans(exp4_exp7_final_model, spec= "Reward_Odour")

Pairwise_exp4_exp7=contrast(Pairwise_exp4_exp7, method = "pairwise")
summary(Pairwise_exp4_exp7)
```

```
##  contrast                    estimate   SE  df z.ratio p.value
##  OdourQSummer - OdourQWinter    -0.99 0.51 Inf  -1.943  0.0520
## 
## Results are given on the log odds ratio (not the response) scale.
```

```
###Merged to one dataset## Note we called the winter betach exp 7 but this was only a working name. 7 = 4b

Merged_4aand4b <- read_excel("C:/Users/LocalAdmin/Documents/ThomasWagnerPhD/PhD/RawData/Associative/OdourLearning/NiceDataStructure/test4and7.xlsx") #plot
Merged4aand4btwo <- read_excel("C:/Users/LocalAdmin/Documents/ThomasWagnerPhD/PhD/RawData/Associative/OdourLearning/NiceDataStructure/exp4_exp7_merged.xlsx") #glmm

Merged_4aand4b_model <-glmer(correct_final ~ Reward_Odour + (1|Colony_ID),
                                  family="binomial",
                                  data= Merged4aand4btwo)
```

```
## boundary (singular) fit: see help('isSingular')
```

```
summary(Merged_4aand4b_model)
```

```
## Generalized linear mixed model fit by maximum likelihood (Laplace
##   Approximation) [glmerMod]
##  Family: binomial  ( logit )
## Formula: correct_final ~ Reward_Odour + (1 | Colony_ID)
##    Data: Merged4aand4btwo
## 
##      AIC      BIC   logLik deviance df.resid 
##    127.3    135.6    -60.6    121.3      117 
## 
## Scaled residuals: 
##     Min      1Q  Median      3Q     Max 
## -2.6458  0.3780  0.3780  0.6202  0.6202 
## 
## Random effects:
##  Groups    Name        Variance Std.Dev.
##  Colony_ID (Intercept) 0        0       
## Number of obs: 120, groups:  Colony_ID, 9
## 
## Fixed effects:
##                          Estimate Std. Error z value Pr(>|z|)    
## (Intercept)                0.9555     0.2631   3.632 0.000282 ***
## Reward_OdourOdourQWinter   0.9904     0.5096   1.943 0.051963 .  
## ---
## Signif. codes:  0 '***' 0.001 '**' 0.01 '*' 0.05 '.' 0.1 ' ' 1
## 
## Correlation of Fixed Effects:
##             (Intr)
## Rwrd_OdrOQW -0.516
## optimizer (Nelder_Mead) convergence code: 0 (OK)
## boundary (singular) fit: see help('isSingular')
```

```
##basic plot merged###

Merged_exp4a_exp4b_plot <- ggplot (Merged_4aand4b, aes(x = Reward_Odour, y = correct_final))+
  scale_y_continuous(expand = c(0, 0)) + # forces X axis to 0, but in this case is overriden by ribbon
  geom_point( alpha = 0) +
  stat_summary(fun.y = "mean", geom = "bar", fill="dodgerblue4") +
  stat_summary(fun.data = "mean_cl_boot", geom="errorbar", width = 0.2) +
  ylab("correct choices") +
  xlab("threatment") +
  theme_bw(26) + theme(axis.line = element_line(colour = "black"),
    panel.grid.major = element_blank(),
    panel.grid.minor = element_blank(),
    panel.border = element_blank(),
    panel.background = element_blank())  +
  geom_abline(intercept = 0.5, slope = 0, color = "black", linetype = 2) +
  coord_cartesian( ylim = c(0.4, 1.05)) # zooms in on the top half
print(Merged_exp4a_exp4b_plot)
```

```
fig4a4bmerged_print <- "fig4a4bmerged_print.pptx"

fig4a4bmerged_final_pp <- dml(ggobj = Merged_exp4a_exp4b_plot)

read_pptx()%>%
add_slide() %>%
ph_with(fig4a4bmerged_final_pp, location = ph_location_fullsize())%>%
print(fig4a4bmerged_print )

ggsave ("fig4a4bmerged_final.png", plot = Merged_exp4a_exp4b_plot, dpi = 300, width = 20, height = 20, units = c("cm"))
```

Ants were able in both experimental sets (summer and winter) to
associate an odour with the reward. Even though, p = 0.052 there is a
strong hint that there might be a experimental set difference especially
in the light of the later following 6th experiment.

#### 5. Analysis, flavour no odour and quinine (summer)

Can ants learn correctly to associate the flavour alone, without
having a runway odour or a punishment visit (quinine), with the reward.
Reward odour is apple 50% or strawberry 50%, Y- maze reward odour 1
vs. odour 2. Data is binomial distributed and colony ID as random
factor.

##### Glmm modeling & pairwise, experiment 5

```
# Glmm

Exp5OnlyFlavourNoQuinine$Reward_odour_flavour<-as.factor(Exp5OnlyFlavourNoQuinine$Reward_odour_flavour)

Exp5OnlyFlavourNoQuinine_m <-glmer(final_correct ~ Reward_odour_flavour + (1|Colony_ID_donor),
                              family="binomial",
                              data= Exp5OnlyFlavourNoQuinine )
```

```
## boundary (singular) fit: see help('isSingular')
```

```
summary(Exp5OnlyFlavourNoQuinine_m)
```

```
## Generalized linear mixed model fit by maximum likelihood (Laplace
##   Approximation) [glmerMod]
##  Family: binomial  ( logit )
## Formula: final_correct ~ Reward_odour_flavour + (1 | Colony_ID_donor)
##    Data: Exp5OnlyFlavourNoQuinine
## 
##      AIC      BIC   logLik deviance df.resid 
##     69.5     75.1    -31.8     63.5       45 
## 
## Scaled residuals: 
##     Min      1Q  Median      3Q     Max 
## -1.2910 -1.2910  0.7746  0.7746  0.7746 
## 
## Random effects:
##  Groups          Name        Variance  Std.Dev. 
##  Colony_ID_donor (Intercept) 2.881e-16 1.697e-08
## Number of obs: 48, groups:  Colony_ID_donor, 2
## 
## Fixed effects:
##                                  Estimate Std. Error z value Pr(>|z|)
## (Intercept)                     5.108e-01  4.216e-01   1.212    0.226
## Reward_odour_flavourStrawberry -1.579e-16  5.963e-01   0.000    1.000
## 
## Correlation of Fixed Effects:
##             (Intr)
## Rwrd_dr_flS -0.707
## optimizer (Nelder_Mead) convergence code: 0 (OK)
## boundary (singular) fit: see help('isSingular')
```

```
# Pairwise

Pairwise5 = emmeans(Exp5OnlyFlavourNoQuinine_m, spec= "Reward_odour_flavour")

Pairwise5=contrast(Pairwise5, method = "pairwise")
summary(Pairwise5)
```

```
##  contrast           estimate    SE  df z.ratio p.value
##  Apple - Strawberry 1.58e-16 0.596 Inf   0.000  1.0000
## 
## Results are given on the log odds ratio (not the response) scale.
```

#### 6. Analysis, flavour no odour and quinine (winter)

We received additional data, this time in winter. This time a
punishment visit (quinine) was added, which is contrary to the 5th
experiment. Reward odour is apple 50% or strawberry 50%, Y- maze reward
odour 1 vs. odour 2. Data is binomial distributed and colony ID as
random factor.

##### Glmm modeling & pairwise, experiment 6

```
# Glmm

Exp6OnlyFlavourQuinineWinter$Reward_Flavour<-as.factor(Exp6OnlyFlavourQuinineWinter$Reward_Flavour)
Exp6OnlyFlavourQuinineWinter_m <-glmer(final_correct ~ Reward_Flavour + (1|Colony_ID),
                                  family="binomial",
                                  data=Exp6OnlyFlavourQuinineWinter )
```

```
## boundary (singular) fit: see help('isSingular')
```

```
summary(Exp6OnlyFlavourQuinineWinter_m)
```

```
## Generalized linear mixed model fit by maximum likelihood (Laplace
##   Approximation) [glmerMod]
##  Family: binomial  ( logit )
## Formula: final_correct ~ Reward_Flavour + (1 | Colony_ID)
##    Data: Exp6OnlyFlavourQuinineWinter
## 
##      AIC      BIC   logLik deviance df.resid 
##     40.3     45.6    -17.1     34.3       41 
## 
## Scaled residuals: 
##     Min      1Q  Median      3Q     Max 
## -3.1623  0.3162  0.3162  0.4714  0.4714 
## 
## Random effects:
##  Groups    Name        Variance Std.Dev.
##  Colony_ID (Intercept) 0        0       
## Number of obs: 44, groups:  Colony_ID, 3
## 
## Fixed effects:
##                          Estimate Std. Error z value Pr(>|z|)   
## (Intercept)                2.3026     0.7416   3.105   0.0019 **
## Reward_FlavourStrawberry  -0.7985     0.9250  -0.863   0.3880   
## ---
## Signif. codes:  0 '***' 0.001 '**' 0.01 '*' 0.05 '.' 0.1 ' ' 1
## 
## Correlation of Fixed Effects:
##             (Intr)
## Rwrd_FlvrSt -0.802
## optimizer (Nelder_Mead) convergence code: 0 (OK)
## boundary (singular) fit: see help('isSingular')
```

```
# Pairwise

Pairwise6 = emmeans(Exp6OnlyFlavourQuinineWinter_m, spec= "Reward_Flavour")

Pairwise6=contrast(Pairwise6, method = "pairwise")
summary(Pairwise6)
```

```
##  contrast           estimate    SE  df z.ratio p.value
##  Apple - Strawberry    0.799 0.925 Inf   0.863  0.3880
## 
## Results are given on the log odds ratio (not the response) scale.
```

Figures of experiment 5 & 6 were merged for a better comparison
in this script. For the manuscript,all short-term experiments were
merged to one big figure. Aesthetic judgement were made via
powerpoint.

##### Figures merged experiment 5 & 6

```
# Basic plot

exp5_exp6_merged_final_plot <- ggplot (exp5_exp6_merged, aes(x = Reward_Odour, y = correct_final))+
  scale_y_continuous(expand = c(0, 0)) + # forces X axis to 0, but in this case is overriden by ribbon
  geom_point( alpha = 0) +
  stat_summary(fun.y = "mean", geom = "bar", fill="dodgerblue4") +
  stat_summary(fun.data = "mean_cl_boot", geom="errorbar", width = 0.2) +
  ylab("correct choices") +
  xlab("threatment") +
  theme_bw(26) + theme(axis.line = element_line(colour = "black"),
    panel.grid.major = element_blank(),
    panel.grid.minor = element_blank(),
    panel.border = element_blank(),
    panel.background = element_blank()) +
  geom_abline(intercept = 0.5, slope = 0, color = "black", linetype = 2) +
  coord_cartesian( ylim = c(0.4, 1.05)) # zooms in on the top half
print(exp5_exp6_merged_final_plot)
```

```
# Powerpoint import

fig_exp5_exp6_final_print <- "fig_exp5_exp6_print.pptx"

fig_exp5_exp6_final_pp <- dml(ggobj = exp5_exp6_merged_final_plot)

read_pptx()%>%
add_slide() %>%
ph_with(fig_exp5_exp6_final_pp, location = ph_location_fullsize())%>%
print(fig_exp5_exp6_final_print)

ggsave ("exp5_exp6_merged_final.png", plot = exp5_exp6_merged_final_plot, dpi = 300, width = 20, height = 20, units = c("cm"))
```

##### Glmm modeling & pairwise comparison experiment 5 vs 6

```
#Glmm


exp5_exp6_final_model <-glmer(correct_final ~ Reward_Odour + (1|Colony_ID),
                                  family="binomial",
                                  data= exp5_exp6_merged)
```

```
## boundary (singular) fit: see help('isSingular')
```

```
summary(exp5_exp6_final_model)
```

```
## Generalized linear mixed model fit by maximum likelihood (Laplace
##   Approximation) [glmerMod]
##  Family: binomial  ( logit )
## Formula: correct_final ~ Reward_Odour + (1 | Colony_ID)
##    Data: exp5_exp6_merged
## 
##      AIC      BIC   logLik deviance df.resid 
##    104.6    112.1    -49.3     98.6       89 
## 
## Scaled residuals: 
##     Min      1Q  Median      3Q     Max 
## -2.5166 -1.2910  0.3974  0.7746  0.7746 
## 
## Random effects:
##  Groups    Name        Variance Std.Dev.
##  Colony_ID (Intercept) 0        0       
## Number of obs: 92, groups:  Colony_ID, 5
## 
## Fixed effects:
##                            Estimate Std. Error z value Pr(>|z|)  
## (Intercept)                  0.5108     0.2981   1.713   0.0866 .
## Reward_OdourFlavourQuinine   1.3350     0.5309   2.515   0.0119 *
## ---
## Signif. codes:  0 '***' 0.001 '**' 0.01 '*' 0.05 '.' 0.1 ' ' 1
## 
## Correlation of Fixed Effects:
##             (Intr)
## Rwrd_OdrFlQ -0.562
## optimizer (Nelder_Mead) convergence code: 0 (OK)
## boundary (singular) fit: see help('isSingular')
```

```
#Pairwise

Pairwise_exp5_exp6_final = emmeans(exp5_exp6_final_model, spec= "Reward_Odour")

Pairwise_exp5_exp6_final=contrast(Pairwise_exp5_exp6_final, method = "pairwise")
summary(Pairwise_exp5_exp6_final)
```

```
##  contrast                          estimate    SE  df z.ratio p.value
##  FlavourNoQuinine - FlavourQuinine    -1.34 0.531 Inf  -2.515  0.0119
## 
## Results are given on the log odds ratio (not the response) scale.
```

Ants were not able to learn in the experimental set 1 (summer) to
associate a flavour alone, without a punishment with a reward. The
results of experimental set 2 (winter) show the opposite. However, the
treatment of the experimental set 2 was slighly different with adding a
punishment visit (quinine). The difference between experiment 5 and 6
could be a season effect or the added punishment visit. Nevertheless,
The results of the 2nd and 3th experiments shift the tendency more to a
seasonal effect.

### Odour-flavour-learning long-term

#### 7. Analysis, flavour and odour no quinine

##### Trained ants

We tested our ants on a Y-maze 6h 24h and 48h after their training.
Every ant was removed from the colony after one test. Therefore every
ant was only tested in one of these 3 time points (6h or 24h or 48). The
aim was to test if ants keep their olfactory memory (odour and flavour
associated with a reward) on a long term view. Reward odour is apple 50%
or strawberry 50%, Y- maze reward odour 1 vs. odour 2. Data is binomial
distributed and colony ID as random factor.

##### Glmm modeling & pairwise, experiment 8

```
# Glmm

Exp8LongTermFlavourOdourNoQuinineTrained$Memory_time<-as.factor(Exp8LongTermFlavourOdourNoQuinineTrained$Memory_time)

summary(Exp8LongTermFlavourOdourNoQuinineTrained)
```

```
##  Colony_Origin      Collection_Date               n_days_starved
##  Length:210         Min.   :2021-06-13 00:00:00   Min.   :4     
##  Class :character   1st Qu.:2021-06-15 00:00:00   1st Qu.:4     
##  Mode  :character   Median :2021-06-16 00:00:00   Median :4     
##                     Mean   :2021-06-16 04:48:00   Mean   :4     
##                     3rd Qu.:2021-06-18 00:00:00   3rd Qu.:4     
##                     Max.   :2021-06-20 00:00:00   Max.   :4     
##                                                                 
##    observer         colony_id_donor colony_id_recipient reward_odour_flavour
##  Length:210         Min.   : 4.0    Min.   : 5.0        Length:210          
##  Class :character   1st Qu.:12.0    1st Qu.: 8.0        Class :character    
##  Mode  :character   Median :14.0    Median :15.0        Mode  :character    
##                     Mean   :13.2    Mean   :13.2                            
##                     3rd Qu.:18.0    3rd Qu.:19.0                            
##                     Max.   :18.0    Max.   :19.0                            
##                                                                             
##  reward_Side        6h_initial_decision 6h_final_decision  24h_initial_decision
##  Length:210         Length:210          Length:210         Length:210          
##  Class :character   Class :character    Class :character   Class :character    
##  Mode  :character   Mode  :character    Mode  :character   Mode  :character    
##                                                                                
##                                                                                
##                                                                                
##                                                                                
##  24h_final_decision 48h_initial_decision 48h_final_decision 6h_correct_initial
##  Length:210         Length:210           Length:210         Min.   :0.0000    
##  Class :character   Class :character     Class :character   1st Qu.:1.0000    
##  Mode  :character   Mode  :character     Mode  :character   Median :1.0000    
##                                                             Mean   :0.8714    
##                                                             3rd Qu.:1.0000    
##                                                             Max.   :1.0000    
##                                                             NA's   :140       
##  6h_correct_final 24h_correct_initial 24h_correct_final 48h_correct_initial
##  Min.   :0.0000   Min.   :0.0000      Min.   :0.0000    Min.   :0.0000     
##  1st Qu.:1.0000   1st Qu.:1.0000      1st Qu.:1.0000    1st Qu.:0.2500     
##  Median :1.0000   Median :1.0000      Median :1.0000    Median :1.0000     
##  Mean   :0.8571   Mean   :0.7714      Mean   :0.8143    Mean   :0.7429     
##  3rd Qu.:1.0000   3rd Qu.:1.0000      3rd Qu.:1.0000    3rd Qu.:1.0000     
##  Max.   :1.0000   Max.   :1.0000      Max.   :1.0000    Max.   :1.0000     
##  NA's   :140      NA's   :140         NA's   :140       NA's   :140        
##  48h_correct_final Memory_time correct_initial  correct_final   
##  Min.   :0.0       24h:70      Min.   :0.0000   Min.   :0.0000  
##  1st Qu.:1.0       48h:70      1st Qu.:1.0000   1st Qu.:1.0000  
##  Median :1.0       6h :70      Median :1.0000   Median :1.0000  
##  Mean   :0.8                   Mean   :0.7952   Mean   :0.8238  
##  3rd Qu.:1.0                   3rd Qu.:1.0000   3rd Qu.:1.0000  
##  Max.   :1.0                   Max.   :1.0000   Max.   :1.0000  
##  NA's   :140
```

```
Exp8LongTermFlavourOdourNoQuinineTrained_m <-glmer(correct_final ~ Memory_time + (1|colony_id_donor),
                                family="binomial",
                                data= Exp8LongTermFlavourOdourNoQuinineTrained)
```

```
## boundary (singular) fit: see help('isSingular')
```

```
summary(Exp8LongTermFlavourOdourNoQuinineTrained_m)
```

```
## Generalized linear mixed model fit by maximum likelihood (Laplace
##   Approximation) [glmerMod]
##  Family: binomial  ( logit )
## Formula: correct_final ~ Memory_time + (1 | colony_id_donor)
##    Data: Exp8LongTermFlavourOdourNoQuinineTrained
## 
##      AIC      BIC   logLik deviance df.resid 
##    202.7    216.1    -97.3    194.7      206 
## 
## Scaled residuals: 
##     Min      1Q  Median      3Q     Max 
## -2.4495  0.4083  0.4776  0.5000  0.5000 
## 
## Random effects:
##  Groups          Name        Variance Std.Dev.
##  colony_id_donor (Intercept) 0        0       
## Number of obs: 210, groups:  colony_id_donor, 4
## 
## Fixed effects:
##                Estimate Std. Error z value Pr(>|z|)    
## (Intercept)     1.47810    0.30735   4.809 1.52e-06 ***
## Memory_time48h -0.09181    0.42866  -0.214    0.830    
## Memory_time6h   0.31366    0.45949   0.683    0.495    
## ---
## Signif. codes:  0 '***' 0.001 '**' 0.01 '*' 0.05 '.' 0.1 ' ' 1
## 
## Correlation of Fixed Effects:
##             (Intr) Mmr_48
## Memry_tm48h -0.717       
## Memory_tm6h -0.669  0.480
## optimizer (Nelder_Mead) convergence code: 0 (OK)
## boundary (singular) fit: see help('isSingular')
```

```
# Pairwise

Pairwise8a = emmeans(Exp8LongTermFlavourOdourNoQuinineTrained_m, spec= "Memory_time")

Pairwise8a=contrast(Pairwise8a, method = "pairwise")
summary(Pairwise8a)
```

```
##  contrast  estimate    SE  df z.ratio p.value
##  24h - 48h   0.0918 0.429 Inf   0.214  0.9750
##  24h - 6h   -0.3137 0.459 Inf  -0.683  0.7736
##  48h - 6h   -0.4055 0.454 Inf  -0.893  0.6444
## 
## Results are given on the log odds ratio (not the response) scale. 
## P value adjustment: tukey method for comparing a family of 3 estimates
```

```
# odour#

Exp8LongTermFlavourOdourNoQuinineTrainedb_m <-glmer(correct_final ~ reward_odour_flavour + (1|colony_id_donor),
                                family="binomial",
                                data= Exp8LongTermFlavourOdourNoQuinineTrained)
summary(Exp8LongTermFlavourOdourNoQuinineTrainedb_m)
```

```
## Generalized linear mixed model fit by maximum likelihood (Laplace
##   Approximation) [glmerMod]
##  Family: binomial  ( logit )
## Formula: correct_final ~ reward_odour_flavour + (1 | colony_id_donor)
##    Data: Exp8LongTermFlavourOdourNoQuinineTrained
## 
##      AIC      BIC   logLik deviance df.resid 
##    200.0    210.1    -97.0    194.0      207 
## 
## Scaled residuals: 
##     Min      1Q  Median      3Q     Max 
## -2.5713  0.3889  0.3936  0.4945  0.5459 
## 
## Random effects:
##  Groups          Name        Variance Std.Dev.
##  colony_id_donor (Intercept) 0.1      0.3163  
## Number of obs: 210, groups:  colony_id_donor, 4
## 
## Fixed effects:
##                                Estimate Std. Error z value Pr(>|z|)    
## (Intercept)                      1.8371     0.4113   4.467 7.94e-06 ***
## reward_odour_flavourStrawberry  -0.6539     0.7065  -0.926    0.355    
## ---
## Signif. codes:  0 '***' 0.001 '**' 0.01 '*' 0.05 '.' 0.1 ' ' 1
## 
## Correlation of Fixed Effects:
##             (Intr)
## rwrd_dr_flS -0.801
```

##### Nestmate

We tested the trained ants´s nestmates which were fed by the trained
ants with the flavoured food reward. The aim was to see if the nestmates
preference can be steered by the forager food intake. Nestmate were
tested after 24h and 48h. An ant was only tested in one of these 2 time
points. Reward odour is apple 50% or strawberry 50%, Y- maze reward
odour 1 vs. odour 2. Data is binomial distributed and colony ID as
random factor.

##### Glmm modeling & pairwise, experiment 8

```
# Glmm

Exp8LongTermFlavourOdourNoQuinineNestmates$Memory_time<-as.factor(Exp8LongTermFlavourOdourNoQuinineNestmates$Memory_time)

Exp8LongTermFlavourOdourNoQuinineNestmates_m <-glmer(correct_final ~ Memory_time + (1|colony_id_donor),
                                              family="binomial",
                                              data=Exp8LongTermFlavourOdourNoQuinineNestmates )
summary(Exp8LongTermFlavourOdourNoQuinineNestmates_m)
```

```
## Generalized linear mixed model fit by maximum likelihood (Laplace
##   Approximation) [glmerMod]
##  Family: binomial  ( logit )
## Formula: correct_final ~ Memory_time + (1 | colony_id_donor)
##    Data: Exp8LongTermFlavourOdourNoQuinineNestmates
## 
##      AIC      BIC   logLik deviance df.resid 
##    190.8    200.1    -92.4    184.8      157 
## 
## Scaled residuals: 
##     Min      1Q  Median      3Q     Max 
## -2.1832 -1.2728  0.5934  0.6065  0.7857 
## 
## Random effects:
##  Groups          Name        Variance Std.Dev.
##  colony_id_donor (Intercept) 0.1207   0.3474  
## Number of obs: 160, groups:  colony_id_donor, 3
## 
## Fixed effects:
##                Estimate Std. Error z value Pr(>|z|)    
## (Intercept)      1.3593     0.3549   3.830 0.000128 ***
## Memory_time48h  -0.5177     0.3625  -1.428 0.153323    
## ---
## Signif. codes:  0 '***' 0.001 '**' 0.01 '*' 0.05 '.' 0.1 ' ' 1
## 
## Correlation of Fixed Effects:
##             (Intr)
## Memry_tm48h -0.579
```

```
# Pairwise

Pairwise8b = emmeans(Exp8LongTermFlavourOdourNoQuinineNestmates_m, spec= "Memory_time")

Pairwise8b=contrast(Pairwise8b, method = "pairwise")
summary(Pairwise8b)
```

```
##  contrast  estimate    SE  df z.ratio p.value
##  24h - 48h    0.518 0.363 Inf   1.428  0.1533
## 
## Results are given on the log odds ratio (not the response) scale.
```

```
# odour#
Exp8LongTermFlavourOdourNoQuinineNestmatesc_m <-glmer(correct_final ~ reward_odour_flavour + (1|colony_id_donor),
                                              family="binomial",
                                              data=Exp8LongTermFlavourOdourNoQuinineNestmates )
```

```
## boundary (singular) fit: see help('isSingular')
```

```
summary(Exp8LongTermFlavourOdourNoQuinineNestmatesc_m)
```

```
## Generalized linear mixed model fit by maximum likelihood (Laplace
##   Approximation) [glmerMod]
##  Family: binomial  ( logit )
## Formula: correct_final ~ reward_odour_flavour + (1 | colony_id_donor)
##    Data: Exp8LongTermFlavourOdourNoQuinineNestmates
## 
##      AIC      BIC   logLik deviance df.resid 
##    188.5    197.7    -91.3    182.5      157 
## 
## Scaled residuals: 
##     Min      1Q  Median      3Q     Max 
## -1.8559 -1.1632  0.5388  0.5388  0.8597 
## 
## Random effects:
##  Groups          Name        Variance Std.Dev.
##  colony_id_donor (Intercept) 0        0       
## Number of obs: 160, groups:  colony_id_donor, 3
## 
## Fixed effects:
##                                Estimate Std. Error z value Pr(>|z|)    
## (Intercept)                      1.2368     0.2186   5.657 1.54e-08 ***
## reward_odour_flavourStrawberry  -0.9345     0.3874  -2.412   0.0159 *  
## ---
## Signif. codes:  0 '***' 0.001 '**' 0.01 '*' 0.05 '.' 0.1 ' ' 1
## 
## Correlation of Fixed Effects:
##             (Intr)
## rwrd_dr_flS -0.564
## optimizer (Nelder_Mead) convergence code: 0 (OK)
## boundary (singular) fit: see help('isSingular')
```

```
#Strawberry was significantly lower after 24h but not after 48h. Therefore lets check if the ants, trained with strawberry, still learnt#


final_straw <- subset (Exp8LongTermFlavourOdourNoQuinineNestmates, reward_odour_flavour=="Strawberry")


final_straw_m <-glmer(correct_final ~ 1 + (1|colony_id_recipient),
                                             family="binomial",
                                             data= Exp8LongTermFlavourOdourNoQuinineNestmates)

summary(final_straw_m)
```

```
## Generalized linear mixed model fit by maximum likelihood (Laplace
##   Approximation) [glmerMod]
##  Family: binomial  ( logit )
## Formula: correct_final ~ 1 + (1 | colony_id_recipient)
##    Data: Exp8LongTermFlavourOdourNoQuinineNestmates
## 
##      AIC      BIC   logLik deviance df.resid 
##    192.1    198.2    -94.0    188.1      158 
## 
## Scaled residuals: 
##     Min      1Q  Median      3Q     Max 
## -1.7798 -1.5629  0.6169  0.6399  0.6399 
## 
## Random effects:
##  Groups              Name        Variance Std.Dev.
##  colony_id_recipient (Intercept) 0.04088  0.2022  
## Number of obs: 160, groups:  colony_id_recipient, 3
## 
## Fixed effects:
##             Estimate Std. Error z value Pr(>|z|)    
## (Intercept)   1.0059     0.2324   4.329  1.5e-05 ***
## ---
## Signif. codes:  0 '***' 0.001 '**' 0.01 '*' 0.05 '.' 0.1 ' ' 1
```

```
### so even though strawberry was significantly weaker after 24h compared to apple the ants still chose significantly more often the reward odour strawberry over apple. Therefore the preference was steered towards the fed flavoured food by the trained ants.
```

##### Figures exp 8, trained ants and nestmates

For the manuscript, all long-term experiments were merged to one big
figure. Aesthetic judgement were made via powerpoint.

```
# trained ants#
Exp8LongTermFlavourOdourNoQuinineTrained$Memory_time <- factor(Exp8LongTermFlavourOdourNoQuinineTrained$Memory_time, levels = c("6h", "24h", "48h"))

fig8anew <- ggplot (Exp8LongTermFlavourOdourNoQuinineTrained, aes(x = Memory_time, y = correct_final))+
  scale_y_continuous(expand = c(0, 0)) + # forces X axis to 0, but in this case is overriden by ribbon
  geom_point( alpha = 0) +
  stat_summary(fun.y = "mean", geom = "bar", fill="dodgerblue4") +
  stat_summary(fun.data = "mean_cl_boot", geom="errorbar", width = 0.2) +
  ylab("Proportion of correct final choices") +
  xlab("Hours after training") +
  theme_bw(26) + theme(axis.line = element_line(colour = "black"),
    panel.grid.major = element_blank(),
    panel.grid.minor = element_blank(),
    panel.border = element_blank(),
    panel.background = element_blank()) +
  geom_abline(intercept = 0.5, slope = 0, color = "black", linetype = 2) +
  coord_cartesian( ylim = c(0.15, 1.05)) # zooms in on the top half
```

```
## Warning: `fun.y` is deprecated. Use `fun` instead.
```

```
print(fig8anew)
```

```
# Powerpoint import

exp8_marked_print <- "exp8_marked_print.pptx"

exp8marked_final_pp <- dml(ggobj = fig8anew)

read_pptx()%>%
add_slide() %>%
ph_with(exp8marked_final_pp, location = ph_location_fullsize())%>%
print(exp8_marked_print)

ggsave ("exp8_marked.png", plot = fig8anew, dpi = 300, width = 20, height = 20, units = c("cm"))

# nestmates

# Basic plot

fig8bnew <- ggplot (Exp8LongTermFlavourOdourNoQuinineNestmates, aes(x = Memory_time, y = correct_final))+
  scale_y_continuous(expand = c(0, 0)) + # forces X axis to 0, but in this case is overriden by ribbon
  geom_point( alpha = 0) +
  stat_summary(fun.y = "mean", geom = "bar", fill="dodgerblue4") +
  stat_summary(fun.data = "mean_cl_boot", geom="errorbar", width = 0.2) +
  ylab("Proportion of correct final choices") +
  xlab("hours after training") +
  theme_bw(26) + theme(axis.line = element_line(colour = "black"),
    panel.grid.major = element_blank(),
    panel.grid.minor = element_blank(),
    panel.border = element_blank(),
    panel.background = element_blank()) +
  geom_abline(intercept = 0.5, slope = 0, color = "black", linetype = 2) +
  coord_cartesian( ylim = c(0.15, 1.05)) # zooms in on the top half
```

```
## Warning: `fun.y` is deprecated. Use `fun` instead.
```

```
print(fig8bnew)
```

```
# Powerpoint import

exp8_unmarked_print <- "exp8_unmarked_print.pptx"

exp8unmarked_final_pp <- dml(ggobj = fig8bnew)

read_pptx()%>%
add_slide() %>%
ph_with(exp8unmarked_final_pp, location = ph_location_fullsize())%>%
print(exp8_unmarked_print)

ggsave ("exp8_unmarked.png", plot = fig8bnew, dpi = 300, width = 20, height = 20, units = c("cm"))
```

Did the nestmate learn significantly weaker than the trained
ants?

##### Glmm modeling & pairwise trained ants vs nestmates, experiment 8

```
# Glmm
Exp8LongTermFlavourOdourNoQuinineTrainedVsNestmates$treatment<-as.factor(Exp8LongTermFlavourOdourNoQuinineTrainedVsNestmates$treatment)

Exp8LongTermFlavourOdourNoQuinineTrainedVsNestmates_m <-glmer(correct_final ~ treatment + (1|colony),
                                                    family="binomial",
                                                    data= Exp8LongTermFlavourOdourNoQuinineTrainedVsNestmates)
summary(Exp8LongTermFlavourOdourNoQuinineTrainedVsNestmates_m)
```

```
## Generalized linear mixed model fit by maximum likelihood (Laplace
##   Approximation) [glmerMod]
##  Family: binomial  ( logit )
## Formula: correct_final ~ treatment + (1 | colony)
##    Data: Exp8LongTermFlavourOdourNoQuinineTrainedVsNestmates
## 
##      AIC      BIC   logLik deviance df.resid 
##    390.0    413.5   -189.0    378.0      364 
## 
## Scaled residuals: 
##     Min      1Q  Median      3Q     Max 
## -3.0351  0.3371  0.4484  0.5680  0.8252 
## 
## Random effects:
##  Groups Name        Variance Std.Dev.
##  colony (Intercept) 0.154    0.3924  
## Number of obs: 370, groups:  colony, 7
## 
## Fixed effects:
##                        Estimate Std. Error z value Pr(>|z|)    
## (Intercept)             1.56862    0.36060   4.350 1.36e-05 ***
## treatment24h untrained -0.18058    0.45402  -0.398    0.691    
## treatment48h trained   -0.09359    0.43143  -0.217    0.828    
## treatment48h untrained -0.70138    0.43676  -1.606    0.108    
## treatment6h trained     0.31892    0.46175   0.691    0.490    
## ---
## Signif. codes:  0 '***' 0.001 '**' 0.01 '*' 0.05 '.' 0.1 ' ' 1
## 
## Correlation of Fixed Effects:
##             (Intr) trt24u trt48t trt48u
## trtmnt24hun -0.675                     
## trtmnt48htr -0.615  0.488              
## trtmnt48hun -0.706  0.668  0.507       
## trtmnt6htrn -0.573  0.458  0.480  0.476
```

```
# Pairwise


Pairwise8tvsn = emmeans(Exp8LongTermFlavourOdourNoQuinineTrainedVsNestmates_m, spec= "treatment")

Pairwise8tvsn=contrast(Pairwise8tvsn, method = "pairwise")
summary(Pairwise8tvsn)
```

```
##  contrast                      estimate    SE  df z.ratio p.value
##  24h trained - 24h untrained     0.1806 0.454 Inf   0.398  0.9947
##  24h trained - 48h trained       0.0936 0.431 Inf   0.217  0.9995
##  24h trained - 48h untrained     0.7014 0.437 Inf   1.606  0.4935
##  24h trained - 6h trained       -0.3189 0.462 Inf  -0.691  0.9586
##  24h untrained - 48h trained    -0.0870 0.449 Inf  -0.194  0.9997
##  24h untrained - 48h untrained   0.5208 0.363 Inf   1.433  0.6062
##  24h untrained - 6h trained     -0.4995 0.477 Inf  -1.047  0.8332
##  48h trained - 48h untrained     0.6078 0.431 Inf   1.410  0.6213
##  48h trained - 6h trained       -0.4125 0.456 Inf  -0.904  0.8954
##  48h untrained - 6h trained     -1.0203 0.461 Inf  -2.215  0.1740
## 
## Results are given on the log odds ratio (not the response) scale. 
## P value adjustment: tukey method for comparing a family of 5 estimates
```

Trained ants were able to associate a flavour and a odour with a food
reward over 6h, 24h and 48h. Even their nestmates preference was steered
after 24h and 48h to the reward odour.

#### 9. Analysis, odour no flavour no quinine

##### Trained ants

We tested our ants on a Y-maze 6h 24h and 48h after their training.
Every ant was removed from the colony after one test. Therefore every
ant was only tested in one of these 3 time points (6h or 24h or 48). The
aim was to test if ants keep their olfactory memory (only odour
associated with a reward) on a long term view. Reward odour is apple 50%
or strawberry 50%, Y- maze reward odour 1 vs. odour 2. Data is binomial
distributed and colony ID as random factor.

##### Glmm modeling & pairwise

```
# Glmm

Exp9LongTermOnlyOdourNoQuinineTrained$Memory_time<-as.factor(Exp9LongTermOnlyOdourNoQuinineTrained$Memory_time)

Exp9LongTermOnlyOdourNoQuinineTrained_m <-glmer(correct_final ~ Memory_time + (1|colony_id_donor),
                                              family="binomial",
                                              data=Exp9LongTermOnlyOdourNoQuinineTrained )
```

```
## boundary (singular) fit: see help('isSingular')
```

```
summary(Exp9LongTermOnlyOdourNoQuinineTrained_m)
```

```
## Generalized linear mixed model fit by maximum likelihood (Laplace
##   Approximation) [glmerMod]
##  Family: binomial  ( logit )
## Formula: correct_final ~ Memory_time + (1 | colony_id_donor)
##    Data: Exp9LongTermOnlyOdourNoQuinineTrained
## 
##      AIC      BIC   logLik deviance df.resid 
##    201.6    214.1    -96.8    193.6      164 
## 
## Scaled residuals: 
##     Min      1Q  Median      3Q     Max 
## -1.9148 -1.4530  0.5222  0.6049  0.6883 
## 
## Random effects:
##  Groups          Name        Variance Std.Dev.
##  colony_id_donor (Intercept) 0        0       
## Number of obs: 168, groups:  colony_id_donor, 4
## 
## Fixed effects:
##                Estimate Std. Error z value Pr(>|z|)    
## (Intercept)      1.2993     0.3257   3.990 6.62e-05 ***
## Memory_time48h  -0.2938     0.4440  -0.662    0.508    
## Memory_time6h   -0.5521     0.4335  -1.273    0.203    
## ---
## Signif. codes:  0 '***' 0.001 '**' 0.01 '*' 0.05 '.' 0.1 ' ' 1
## 
## Correlation of Fixed Effects:
##             (Intr) Mmr_48
## Memry_tm48h -0.734       
## Memory_tm6h -0.751  0.551
## optimizer (Nelder_Mead) convergence code: 0 (OK)
## boundary (singular) fit: see help('isSingular')
```

```
# Pairwise


Pairwise9a = emmeans(Exp9LongTermOnlyOdourNoQuinineTrained_m, spec= "Memory_time")

Pairwise9a=contrast(Pairwise9a, method = "pairwise")
summary(Pairwise9a)
```

```
##  contrast  estimate    SE  df z.ratio p.value
##  24h - 48h    0.294 0.444 Inf   0.662  0.7857
##  24h - 6h     0.552 0.434 Inf   1.273  0.4102
##  48h - 6h     0.258 0.416 Inf   0.621  0.8085
## 
## Results are given on the log odds ratio (not the response) scale. 
## P value adjustment: tukey method for comparing a family of 3 estimates
```

```
# odour#

Exp9LongTermOnlyOdourNoQuinineTrainedb_m <-glmer(correct_final ~ reward_odour_flavour + (1|colony_id_donor),
                                              family="binomial",
                                              data=Exp9LongTermOnlyOdourNoQuinineTrained )
```

```
## boundary (singular) fit: see help('isSingular')
```

```
summary(Exp9LongTermOnlyOdourNoQuinineTrainedb_m)
```

```
## Generalized linear mixed model fit by maximum likelihood (Laplace
##   Approximation) [glmerMod]
##  Family: binomial  ( logit )
## Formula: correct_final ~ reward_odour_flavour + (1 | colony_id_donor)
##    Data: Exp9LongTermOnlyOdourNoQuinineTrained
## 
##      AIC      BIC   logLik deviance df.resid 
##    201.2    210.6    -97.6    195.2      165 
## 
## Scaled residuals: 
##     Min      1Q  Median      3Q     Max 
## -1.6787 -1.6285  0.5957  0.6140  0.6140 
## 
## Random effects:
##  Groups          Name        Variance  Std.Dev. 
##  colony_id_donor (Intercept) 1.461e-14 1.209e-07
## Number of obs: 168, groups:  colony_id_donor, 4
## 
## Fixed effects:
##                                Estimate Std. Error z value Pr(>|z|)    
## (Intercept)                     1.03609    0.24816   4.175 2.98e-05 ***
## reward_odour_flavourStrawberry -0.06071    0.34850  -0.174    0.862    
## ---
## Signif. codes:  0 '***' 0.001 '**' 0.01 '*' 0.05 '.' 0.1 ' ' 1
## 
## Correlation of Fixed Effects:
##             (Intr)
## rwrd_dr_flS -0.712
## optimizer (Nelder_Mead) convergence code: 0 (OK)
## boundary (singular) fit: see help('isSingular')
```

##### Nestmate

We tested the trained ants´s nestmates which were fed by the trained
ants with the food reward but this time without added flavour. The aim
was to see if the nestmates preference can be steered by the forager
food intake even when their is no flavour cue around them. Nestmate were
tested after 24h and 48h. An ant was only tested in one of these 2 time
points. Reward odour is apple 50% or strawberry 50%, Y- maze reward
odour 1 vs. odour 2. Data is binomial distributed and colony ID as
random factor.

##### Glmm modeling & pairwise, experiment 9

```
# Glmm

Exp9LongTermOnlyOdourNoQuinineNestmates$Memory_time<-as.factor(Exp9LongTermOnlyOdourNoQuinineNestmates$Memory_time)


Exp9LongTermOnlyOdourNoQuinineNestmates_m <-glmer(correct_final ~ Memory_time + (1|colony_id_donor),
                                                family="binomial",
                                                data= Exp9LongTermOnlyOdourNoQuinineNestmates)
```

```
## boundary (singular) fit: see help('isSingular')
```

```
summary(Exp9LongTermOnlyOdourNoQuinineNestmates_m)
```

```
## Generalized linear mixed model fit by maximum likelihood (Laplace
##   Approximation) [glmerMod]
##  Family: binomial  ( logit )
## Formula: correct_final ~ Memory_time + (1 | colony_id_donor)
##    Data: Exp9LongTermOnlyOdourNoQuinineNestmates
## 
##      AIC      BIC   logLik deviance df.resid 
##    211.4    220.6   -102.7    205.4      157 
## 
## Scaled residuals: 
##     Min      1Q  Median      3Q     Max 
## -0.8165 -0.8165 -0.6546  1.2247  1.5275 
## 
## Random effects:
##  Groups          Name        Variance Std.Dev.
##  colony_id_donor (Intercept) 0        0       
## Number of obs: 160, groups:  colony_id_donor, 4
## 
## Fixed effects:
##                Estimate Std. Error z value Pr(>|z|)  
## (Intercept)     -0.4055     0.2282  -1.777   0.0756 .
## Memory_time48h  -0.4418     0.3341  -1.323   0.1860  
## ---
## Signif. codes:  0 '***' 0.001 '**' 0.01 '*' 0.05 '.' 0.1 ' ' 1
## 
## Correlation of Fixed Effects:
##             (Intr)
## Memry_tm48h -0.683
## optimizer (Nelder_Mead) convergence code: 0 (OK)
## boundary (singular) fit: see help('isSingular')
```

```
# Pairwise

Pairwise9b = emmeans(Exp9LongTermOnlyOdourNoQuinineNestmates_m, spec= "Memory_time")

Pairwise9b=contrast(Pairwise9b, method = "pairwise")
summary(Pairwise9b)
```

```
##  contrast  estimate    SE  df z.ratio p.value
##  24h - 48h    0.442 0.334 Inf   1.323  0.1860
## 
## Results are given on the log odds ratio (not the response) scale.
```

```
## odour #
Exp9LongTermOnlyOdourNoQuinineNestmatesc_m <-glmer(correct_final ~ reward_odour_flavour + (1|colony_id_donor),
                                                family="binomial",
                                                data= Exp9LongTermOnlyOdourNoQuinineNestmates)
```

```
## boundary (singular) fit: see help('isSingular')
```

```
summary(Exp9LongTermOnlyOdourNoQuinineNestmatesc_m)
```

```
## Generalized linear mixed model fit by maximum likelihood (Laplace
##   Approximation) [glmerMod]
##  Family: binomial  ( logit )
## Formula: correct_final ~ reward_odour_flavour + (1 | colony_id_donor)
##    Data: Exp9LongTermOnlyOdourNoQuinineNestmates
## 
##      AIC      BIC   logLik deviance df.resid 
##    211.4    220.6   -102.7    205.4      157 
## 
## Scaled residuals: 
##     Min      1Q  Median      3Q     Max 
## -0.8165 -0.8165 -0.6546  1.2247  1.5275 
## 
## Random effects:
##  Groups          Name        Variance Std.Dev.
##  colony_id_donor (Intercept) 0        0       
## Number of obs: 160, groups:  colony_id_donor, 4
## 
## Fixed effects:
##                                Estimate Std. Error z value Pr(>|z|)  
## (Intercept)                     -0.4055     0.2282  -1.777   0.0756 .
## reward_odour_flavourStrawberry  -0.4418     0.3341  -1.323   0.1860  
## ---
## Signif. codes:  0 '***' 0.001 '**' 0.01 '*' 0.05 '.' 0.1 ' ' 1
## 
## Correlation of Fixed Effects:
##             (Intr)
## rwrd_dr_flS -0.683
## optimizer (Nelder_Mead) convergence code: 0 (OK)
## boundary (singular) fit: see help('isSingular')
```

##### Figures exp 9, trained ants and nestmates

```
# Basic plot

Exp9LongTermOnlyOdourNoQuinineTrained$Memory_time <- factor(Exp9LongTermOnlyOdourNoQuinineTrained$Memory_time, levels = c("6h", "24h", "48h"))

fig9anew <- ggplot (Exp9LongTermOnlyOdourNoQuinineTrained, aes(x = Memory_time, y = correct_final))+
  scale_y_continuous(expand = c(0, 0)) + # forces X axis to 0, but in this case is overriden by ribbon
  geom_point( alpha = 0) +
  stat_summary(fun.y = "mean", geom = "bar", fill="dodgerblue4") +
  stat_summary(fun.data = "mean_cl_boot", geom="errorbar", width = 0.2) +
  ylab("Proportion of correct final choices") +
  xlab("Hours after training") +
  theme_bw(26) + theme(axis.line = element_line(colour = "black"),
    panel.grid.major = element_blank(),
    panel.grid.minor = element_blank(),
    panel.border = element_blank(),
    panel.background = element_blank()) +
  geom_abline(intercept = 0.5, slope = 0, color = "black", linetype = 2) +
  coord_cartesian( ylim = c(0.15, 1.05)) # zooms in on the top half
```

```
## Warning: `fun.y` is deprecated. Use `fun` instead.
```

```
print(fig9anew)
```

```
# Powerpoint import

exp9_marked_print <- "exp9_marked_print.pptx"

exp9marked_final_pp <- dml(ggobj = fig9anew)

read_pptx()%>%
add_slide() %>%
ph_with(exp9marked_final_pp, location = ph_location_fullsize())%>%
print(exp9_marked_print)

ggsave ("exp9_marked.png", plot = fig9anew, dpi = 300, width = 20, height = 20, units = c("cm"))

# Nestmates

# Basic plot

Exp9LongTermOnlyOdourNoQuinineNestmates$Memory_time <- factor(Exp9LongTermOnlyOdourNoQuinineNestmates$Memory_time, levels = c("6h", "24h", "48h"))

fig9bnew <- ggplot (Exp9LongTermOnlyOdourNoQuinineNestmates, aes(x = Memory_time, y = correct_final))+
  scale_y_continuous(expand = c(0, 0)) + # forces X axis to 0, but in this case is overriden by ribbon
  geom_point( alpha = 0) +
  stat_summary(fun.y = "mean", geom = "bar", fill="dodgerblue4") +
  stat_summary(fun.data = "mean_cl_boot", geom="errorbar", width = 0.2) +
  ylab("Proporton of correct final choices") +
  xlab("Hours after training") +
  theme_bw(26) + theme(axis.line = element_line(colour = "black"),
    panel.grid.major = element_blank(),
    panel.grid.minor = element_blank(),
    panel.border = element_blank(),
    panel.background = element_blank()) +
  geom_abline(intercept = 0.5, slope = 0, color = "black", linetype = 2) +
  coord_cartesian( ylim = c(0.15, 1.05)) # zooms in on the top half
```

```
## Warning: `fun.y` is deprecated. Use `fun` instead.
```

```
print(fig9bnew)
```

```
# Powerpoint import

exp9_unmarked_print <- "exp9_unmarked_print.pptx"

exp9unmarked_final_pp <- dml(ggobj = fig9bnew)

read_pptx()%>%
add_slide() %>%
ph_with(exp9unmarked_final_pp, location = ph_location_fullsize())%>%
print(exp9_unmarked_print)

ggsave ("exp9_unmarked.png", plot = fig9bnew, dpi = 300, width = 20, height = 20, units = c("cm"))
```

Did the nestmate learned significantly weaker than the trained
ants?

##### Glmm modeling & pairwise trained ants vs nestmates, experiment 9

```
# Glmm


Exp9LongTermFlavourOdourNoQuinineTrainedVsNestmates$treatment<-as.factor(Exp9LongTermFlavourOdourNoQuinineTrainedVsNestmates$treatment)


Exp9LongTermFlavourOdourNoQuinineTrainedVsNestmates_m <-glmer(correct_final ~ treatment + (1|colony),
                                  family="binomial",
                                  data=Exp9LongTermFlavourOdourNoQuinineTrainedVsNestmates )
```

```
## boundary (singular) fit: see help('isSingular')
```

```
summary(Exp9LongTermFlavourOdourNoQuinineTrainedVsNestmates_m)
```

```
## Generalized linear mixed model fit by maximum likelihood (Laplace
##   Approximation) [glmerMod]
##  Family: binomial  ( logit )
## Formula: correct_final ~ treatment + (1 | colony)
##    Data: Exp9LongTermFlavourOdourNoQuinineTrainedVsNestmates
## 
##      AIC      BIC   logLik deviance df.resid 
##    411.0    433.8   -199.5    399.0      322 
## 
## Scaled residuals: 
##     Min      1Q  Median      3Q     Max 
## -1.9148 -0.8165  0.5222  0.6883  1.5275 
## 
## Random effects:
##  Groups Name        Variance  Std.Dev. 
##  colony (Intercept) 2.075e-14 1.441e-07
## Number of obs: 328, groups:  colony, 7
## 
## Fixed effects:
##                        Estimate Std. Error z value Pr(>|z|)    
## (Intercept)              1.2993     0.3257   3.990 6.62e-05 ***
## treatment24h untrained  -1.7047     0.3977  -4.287 1.81e-05 ***
## treatment48h trained    -0.2938     0.4440  -0.662    0.508    
## treatment48h untrained  -2.1466     0.4069  -5.275 1.33e-07 ***
## treatment6h trained     -0.5521     0.4335  -1.273    0.203    
## ---
## Signif. codes:  0 '***' 0.001 '**' 0.01 '*' 0.05 '.' 0.1 ' ' 1
## 
## Correlation of Fixed Effects:
##             (Intr) trt24u trt48t trt48u
## trtmnt24hun -0.819                     
## trtmnt48htr -0.734  0.601              
## trtmnt48hun -0.800  0.655  0.587       
## trtmnt6htrn -0.751  0.615  0.551  0.601
## optimizer (Nelder_Mead) convergence code: 0 (OK)
## boundary (singular) fit: see help('isSingular')
```

```
# Pairwise

Pairwise9tvsn = emmeans(Exp9LongTermFlavourOdourNoQuinineTrainedVsNestmates_m, spec= "treatment")


Pairwise9tvsn=contrast(Pairwise9tvsn, method = "pairwise")
summary(Pairwise9tvsn)
```

```
##  contrast                      estimate    SE  df z.ratio p.value
##  24h trained - 24h untrained      1.705 0.398 Inf   4.287  0.0002
##  24h trained - 48h trained        0.294 0.444 Inf   0.662  0.9645
##  24h trained - 48h untrained      2.147 0.407 Inf   5.275  <.0001
##  24h trained - 6h trained         0.552 0.434 Inf   1.273  0.7076
##  24h untrained - 48h trained     -1.411 0.378 Inf  -3.729  0.0018
##  24h untrained - 48h untrained    0.442 0.334 Inf   1.323  0.6771
##  24h untrained - 6h trained      -1.153 0.366 Inf  -3.149  0.0141
##  48h trained - 48h untrained      1.853 0.388 Inf   4.775  <.0001
##  48h trained - 6h trained         0.258 0.416 Inf   0.621  0.9718
##  48h untrained - 6h trained      -1.595 0.376 Inf  -4.240  0.0002
## 
## Results are given on the log odds ratio (not the response) scale. 
## P value adjustment: tukey method for comparing a family of 5 estimates
```

Trained ants were able to associate only one cue (odour) with the
food reward after 6h, 24h and 48h. Unlike in the 8th experiment,
nestmates did not show a preference for the odour leading to the reward
anymore. Probably because of the lack of flavour cue (compare with
experiment 8).

### Analysis Summary

- Ants can rapidly form lang lasting (up to 48h) associations between
  two cues (odour and flavour) or only cue (odour or flavour) with a food
  reward reward.
- A punishment visit (quinine) seems to be not needed to form strong
  associative memories.
- Even nestmates`s preference can be steered through the foragers food
  intake as long the food is flavoured.
- There might be a season effect in learning.

### Package info

```
## R version 4.2.1 (2022-06-23 ucrt)
## Platform: x86_64-w64-mingw32/x64 (64-bit)
## Running under: Windows 10 x64 (build 19044)
## 
## Matrix products: default
## 
## locale:
## [1] LC_COLLATE=German_Germany.utf8  LC_CTYPE=German_Germany.utf8   
## [3] LC_MONETARY=German_Germany.utf8 LC_NUMERIC=C                   
## [5] LC_TIME=German_Germany.utf8    
## 
## attached base packages:
## [1] grid      stats     graphics  grDevices utils     datasets  methods  
## [8] base     
## 
## other attached packages:
##  [1] gdtools_0.2.4     cowplot_1.1.1     eoffice_0.2.1     Hmisc_4.7-1      
##  [5] Formula_1.2-4     rvg_0.2.5         officer_0.4.3     EnvStats_2.7.0   
##  [9] lattice_0.20-45   scales_1.2.1      gridExtra_2.3     wesanderson_0.3.6
## [13] readxl_1.4.1      DHARMa_0.4.5      multcomp_1.4-20   TH.data_1.1-1    
## [17] MASS_7.3-57       survival_3.3-1    mvtnorm_1.1-3     emmeans_1.8.0    
## [21] forcats_0.5.2     stringr_1.4.1     dplyr_1.0.9       purrr_0.3.4      
## [25] readr_2.1.2       tidyr_1.2.0       tibble_3.1.8      tidyverse_1.3.2  
## [29] nlme_3.1-157      lme4_1.1-30       Matrix_1.4-1      ggpubr_0.4.0     
## [33] gplots_3.1.3      ggplot2_3.3.6    
## 
## loaded via a namespace (and not attached):
##   [1] uuid_1.1-0          backports_1.4.1     R.devices_2.17.1   
##   [4] systemfonts_1.0.4   lazyeval_0.2.2      splines_4.2.1      
##   [7] gap.datasets_0.0.5  digest_0.6.29       yulab.utils_0.0.5  
##  [10] htmltools_0.5.3     magick_2.7.3        fansi_1.0.3        
##  [13] magrittr_2.0.3      checkmate_2.1.0     googlesheets4_1.0.1
##  [16] cluster_2.1.3       tzdb_0.3.0          modelr_0.1.9       
##  [19] R.utils_2.12.0      sandwich_3.0-2      jpeg_0.1-9         
##  [22] colorspace_2.0-3    rvest_1.0.3         haven_2.5.1        
##  [25] xfun_0.32           crayon_1.5.1        jsonlite_1.8.0     
##  [28] zoo_1.8-10          glue_1.6.2          gtable_0.3.0       
##  [31] gargle_1.2.0        car_3.1-0           abind_1.4-5        
##  [34] DBI_1.1.3           rstatix_0.7.0       Rcpp_1.0.9         
##  [37] viridisLite_0.4.1   xtable_1.8-4        htmlTable_2.4.1    
##  [40] gridGraphics_0.5-1  foreign_0.8-82      htmlwidgets_1.5.4  
##  [43] httr_1.4.4          RColorBrewer_1.1-3  ellipsis_0.3.2     
##  [46] farver_2.1.1        pkgconfig_2.0.3     R.methodsS3_1.8.2  
##  [49] nnet_7.3-17         sass_0.4.2          dbplyr_2.2.1       
##  [52] deldir_1.0-6        utf8_1.2.2          labeling_0.4.2     
##  [55] ggplotify_0.1.0     tidyselect_1.1.2    rlang_1.0.4        
##  [58] munsell_0.5.0       cellranger_1.1.0    tools_4.2.1        
##  [61] cachem_1.0.6        cli_3.3.0           generics_0.1.3     
##  [64] devEMF_4.1          broom_1.0.0         evaluate_0.16      
##  [67] fastmap_1.1.0       yaml_2.3.5          knitr_1.39         
##  [70] fs_1.5.2            zip_2.2.0           caTools_1.18.2     
##  [73] R.oo_1.25.0         xml2_1.3.3          gap_1.2.3-6        
##  [76] compiler_4.2.1      rstudioapi_0.14     plotly_4.10.0      
##  [79] png_0.1-7           ggsignif_0.6.3      reprex_2.0.2       
##  [82] bslib_0.4.0         stringi_1.7.8       highr_0.9          
##  [85] nloptr_2.0.3        vctrs_0.4.1         pillar_1.8.1       
##  [88] lifecycle_1.0.1     jquerylib_0.1.4     estimability_1.4.1 
##  [91] data.table_1.14.2   bitops_1.0-7        flextable_0.7.3    
##  [94] R6_2.5.1            latticeExtra_0.6-30 KernSmooth_2.23-20 
##  [97] codetools_0.2-18    boot_1.3-28         gtools_3.9.3       
## [100] assertthat_0.2.1    withr_2.5.0         hms_1.1.2          
## [103] rpart_4.1.16        minqa_1.2.4         rmarkdown_2.15     
## [106] carData_3.0-5       googledrive_2.0.0   lubridate_1.8.0    
## [109] base64enc_0.1-3     interp_1.1-3
```
