## Supplements 3 - Pilot-Study and Side-learning pairwise tabletrash for "A systematic examination of learning in the invasive ant *Linepithema humile* reveals very rapid development of short and long-term memory"

1. **Pilot-experiment - Odour preference test**

Ants were tested on a Y-maze for their odour preference. One arm was covered with an apple-scented paper overlay, while the other arm was covered with a strawberry-scented overlay. Naïve ants were allowed to freely run up a bridge and onto the Y-maze. The number of ants reaching each end of the maze was counted, and the ant removed.

Our tested ants (n = 158) showed a small but significant preference for strawberry odour (x = 92 Strawberry) over apple odour (binomial test: p < 0.04637).

2. **Experiment 1, Side-learning**

Table S1: 1. Experiment: Side-learning, pairwise comparison (visits).

| **Pairwise comparison** | **SE** | **z. ratio** | **p-value** |
| --- | --- | --- | --- |
| 2 vs. 3 | 0.563 | -2.223 | 0.1169 |
| 2 vs. 4 | 0.559 | -2.139 | 0.1408 |
| **2 vs. 5** | **0.613** | **-2.589** | **0.0474*** |
| 3 vs. 4 | 0.612 | 0.092 | 0.9997 |
| 3 vs. 5 | 0.657 | -0.511 | 0.9566 |
| 4 vs. 5 | 0.657 | -0.596 | 0.9334 |
