## Supplements 4 - Detailed results with statistics AB for "A systematic examination of learning in the invasive ant *Linepithema humile* reveals very rapid development of short and long-term memory"

Supplement 4 – results supplement

This supplement provides a detailed account of the results summarised in the main manuscript. For the complete statistical analysis output, see supplement 1.

### Odour association: Short-term memory

##### Experiment 2 - Short-term olfactory-memory: scented runway and flavoured reward versus punishment

This experiment tested if ants learn to associate the sucrose reward with both environmental cues, a food flavour and the corresponding runway odour (apple or strawberry). 84% (61/72) of the ants chose the food-associated odour arm (GLMM, z-ratio = 3.375, p < 0.001, see fig. 3). The specific odour rewarded did not significantly affect choice accuracy (apple 80% and strawberry 88%, z-ratio = -0.972, p = 0.3312).

##### Experiment 3 – Short-term olfactory-memory: scented runway and flavoured reward versus neutral stimulus

This experiment tested whether a reward alone, and no punishment (quinine), results in an equally high proportion of choices for the food-associated odour. 85% (41/48) of the ants chose food-associated odour arm (GLMM, z-ratio = 2.938, p = 0.003, see fig. 3). Again, the specific odour rewarded did not significantly affect choice accuracy (apple 83% and strawberry 87%, z-ratio = -0.408, p = 0.683).

##### Experiment 4 - Short-term olfactory-memory: scented runway and unflavoured reward versus punishment

This experiment tested if Argentine ants can associate a runway odour (apple or strawberry) with an unflavoured reward. In the summer batch, 72% (52/72) of the ants chose the side of the Y-maze with a scent associated with a reward (GLMM, z-ratio = 2.568, p = 0.010). Ants performed identically on both rewarded odours (72% correct decisions for both odours, z-ratio = 0.00, p = 1.0000, see fig. 3). Merging both batches, showed that 78.5% (94/120) of the ants chose food-associated odour arm (GLMM, z-ratio = 3.632, p < 0.001, see fig. 3)

##### Experiment 5 - Short-term olfactory-memory: unscented runway and flavoured reward versus neutral stimulus

This experiment was run to test the ant´s learning ability without negative reinforcement and using only one cue, the flavoured food. 62.5% (30/48) of ants chose the food-associated odour arm, which does not differ significantly from random choice (GLMM, z-ratio = 1.212, p = 0.226, see fig. 3). Again, there was no significant difference between performance with the two odours (62.5% correct decisions for both odours, z-ratio = 0.000, p = 1.000).

##### Experiment 6 - Short-term olfactory-memory: unscented runway and flavoured reward versus punishment

This experiment was similar as the 5^th^ experiment except that a quinine visit was added and the experiment was conducted in winter. Again, the purpose of this study was to test for the need of a punishment visit. Unlike in summer (experiment 5), the winter experiment (experiment 6) showed that ants were able to learn to associate just one cue (flavour) with a sucrose reward. 86% (38/44) of the ants chose the side of the Y-maze with a scent associated with reward (GLMM, z-ratio = 2.721, p = 0.0065, see fig. 3. There was no significant difference between the two odours (apple 81% and strawberry 90%, z-ratio = -0.863, p = 0.388). A pairwise comparison between the 5^th^ and the 6^th^ experiment showed that ants learned significantly better in the 6^th^ experiment than in the 5^th^ (emmeans, pairwise: z-ratio = -2.515, p = 0.011, see figure 3)

##### Experiment 7 - Short-term olfactory-memory: scented runway and unflavoured reward versus punishment

Like in the summer experiment (4^th^), the ants formed in the winter again a strong association between just one cue (odour) and the sucrose reward. 87.5% (42/48) of the ants chose the side of the Y-maze with a scent associated with reward (GLMM, z-ratio = 2.656, p = 0.007, see fig. 3). Again, the specific odour rewarded did not significantly affect choice accuracy (apple 79% and strawberry 95%, z-ratio = -1.582, p = 0.1138). A pairwise comparison between the 4^th^ and the 7^th^ experiment, thus seasons, showed that ants tend to learn better in the winter (emmeans, pairwise: z-ratio = -1.943, p = 0.052, see figure 3).

###### The roles of runway odour and food flavour in learning

In order to characterise the relative roles of runway odour and food flavour in driving learning, we compared the behaviour of ants in experiments 2 (runway odour and food flavour during training) with the results from experiment 4 (only runway odour) and experiment 5 (only food flavour). Note, however, that experiment 5 did not include a negative reinforcement, while experiments 2 and 4 did. We consider this appropriate, since negative reinforcement did not improve learning (compare experiments 2 & 3 and see possible season effect).

Combined runway odour and flavoured food led to a significantly higher proportion of correct choices than only having flavoured food (z-ratio = -2.714, p = 0.018). However, runway odour alone was not significantly weaker than the combined cues (z-ratio = -1.118, p = 0.502). Runway odour alone did not result in significantly more correct choices than food flavour alone (z-ratio = 1.803, p = 0.168).

### Odour association: Long-term memory

##### Experiment 8 - Long-term olfactory-memory: scented runway and flavoured reward versus neutral stimulus

###### Trained ants and their ability to build a long-term association

Ants trained on combined runway odour and food flavour significantly preferred the odour-associated Y-maze arm after 6 hours, 24 hours and 48 hours (GLMM, z-ratio = 4.204, p < 0.0001, see fig. 4a), with 85% (60/70), 81% (57/70), and 80% (56/70) correct choices after 6, 24, and 48 hours respectively (6 hours: GLMM, z-ratio = 5.246 , p < 0.0001, 24 hours: GLMM, z-ratio = 4.809, p < 0.0001 , 48 hours: GLMM, z-ratio = 8.515, p < 0.0001).

###### Odour preference in untrained ants housed with flavoured-food fed nestmates (trained)

Untrained ants housed with ants trained with flavoured food showed a strong preference for that food flavour after 24 and 48 hours (z-ratio = 3.830, p < 0.0001, see fig. 4a). 77% (62/80) and 67% (54/80) of ants chose food-associated odour after 24 and 48 hours respectively (24 hours: GLMM, z-ratio = 4.619, p < 0.0001, 48 hours: GLMM, n = 80, z-ratio = 2.511, p = 0.012).

##### Experiment 9 - Long-term olfactory-memory: scented runway and unflavoured reward versus neutral stimulus

This experiment was run to quantify long term learning without access to informative flavoured food, since this could refresh the ants’ memories.

###### Trained ants and their ability to build a long-term association with unflavoured food

Ants trained with runway odour and unflavoured food nonetheless significantly preferred the odour-associated Y-maze arm after 6, 24 and 48 hours (GLMM, z-ratio = 3.990, p < 0.0001, see fig. 4b), with 67% (38/56), 78% (44/56), 73% (41/56) correct choices after 6, 24, and 48 hours respectively (6 hours: GLMM, z-ratio = 2.611 , p = 0.009, 24 hours: GLMM, z-ratio = 3.990, p < 0.0001 , 48 hours: GLMM, z-ratio = 3.332, p < 0.0001).

###### Odour preference in untrained ants housed with unflavoured-food fed nestmates (trained)

Unlike naïve ants housed with nestmates trained on flavoured food, naïve ants housed with nestmates trained on unflavoured food showed no preference for the food-associated odour (z-ratio = -1.777, p = 0.075, see fig. 4b). 40% (32/80) and 30% (24/80) of ants chose the food-odour covered Y-maze arm after 24 and 48 hours, respectively. Indeed, while the choice of naïve ants after 24 hours did not differ from chance (GLMM, z-ratio = -1.381, p = 0.268), after 48 hours ants significantly avoided the scented arm (GLMM, z-ratio = -3.473, p < 0.0001).

###### The effect of food flavour presence on the long-term memory of trained ants

Here we compared how well the Argentine ants performed on the long-term memory test (6h, 24h, 48h) when they had a flavoured reward vs. an unflavoured reward. Do they form a significantly better memories when they had a flavoured reward?

There was no significant difference between trained ants who had an unflavoured reward treatment (Experiment 9) and those trained on flavoured food (experiment 8) after 6 hours, 24 hours and 48 hours (6 hours: GLMM, n =56[exp.7] and n=70[exp.8] , z-ratio = 2.344 , p = 0.176 ,24 hours: GLMM, n =56[exp.7] and n=70[exp.8] , z-ratio = 0.399 , p = 0.998, 48 hours: GLMM, n =56[exp.7] and n=70[exp.8] , z-ratio = 0.897, p = 0.947).

However, untrained ants who were treated with an unflavoured reward (9. Experiment) performed significantly worse than ants with a flavoured reward treatment (8. Experiment) after 24 hours and 48 hours (24 hours: GLMM, n =80[exp.7] and n=80[exp.8], z-ratio = 4.709, p < 0.001, 48 hours: GLMM, n =80[exp.7] and n=80[exp.8], z-ratio = 4.631, p < 0.001).
